# Atlas of Subcellular RNA Localization Revealed by APEX-seq

**DOI:** 10.1101/454470

**Authors:** Furqan M. Fazal, Shuo Han, Pornchai Kaewsapsak, Kevin R. Parker, Jin Xu, Alistair N. Boettiger, Howard Y. Chang, Alice Y. Ting

## Abstract

We introduce APEX-seq, a method for RNA sequencing based on spatial proximity to the peroxidase enzyme APEX2. APEX-seq in nine distinct subcellular locales produced a nanometer-resolution spatial map of the human transcriptome, revealing extensive and exquisite patterns of localization for diverse RNA classes and transcript isoforms. We uncover a radial organization of the nuclear transcriptome, which is gated at the inner surface of the nuclear pore for cytoplasmic export of processed transcripts. We identify two distinct pathways of messenger RNA localization to mitochondria, each associated with specific sets of transcripts for building complementary macromolecular machines within the organelle. APEX-seq should be widely applicable to many systems, enabling comprehensive investigations of the spatial transcriptome.

## INTRODUCTION

The subcellular localization of RNA is intimately tied to its function(Buxbaum et al., 2015). Asymmetrically-distributed RNAs underlie organismal development, local protein translation, and the 3D organization of chromatin. Where an RNA is located within the cell likely determines whether it will be stored, processed(Buxbaum et al., 2015; Chin and Lecuyer, 2017), translated(Berkovits and Mayr, 2015), or degraded(Fasken and Corbett, 2009). Consequently, a major goal of biological research is to elucidate the subcellular organization of RNAs, i.e., produce a spatial map of the transcriptome.

While many methods have been developed to study RNA localization(Weil et al., 2010), only a few have been applied on a transcriptome-wide scale. One approach is biochemical fractionation to enrich specific organelles, followed by RNA sequencing (“fractionation-seq”). This method has been applied to transcriptome analysis of mitochondria(Mercer et al., 2011), endoplasmic reticulum (ER)(Reid and Nicchitta, 2012), nucleus and cytosol(Djebali et al., 2012), and stress granules(Khong et al., 2017). However, a major limitation of fractionation-seq is that it can only be applied to organelles that are possible to purify. Many subcellular compartments, such as the nuclear lamina, outer mitochondrial membrane, and pre- and post-synaptic termini, are impossible to purify but hold great interest. Even for compartments that *can* be enriched, such as mitochondria, current protocols fail to remove many contaminants (such as co-purifying RNAs from ER, nuclear, peroxisome, or plasma membranes and cytosolic ribosomes(Sadowski et al., 2008) – leading to high false positive rates) and may lose material of interest (such as RNAs weakly associated with the outer mitochondrial membrane(Lesnik et al., 2015) - leading to false negatives).

RNA localization can also be directly visualized by microscopy(Bertrand et al., 1998; Femino et al., 1998), and the MERFISH(Chen et al., 2015b) and SeqFISH(Shah et al., 2016)) techniques have recently been pioneered for imaging thousands of cellular RNAs at once using barcoded oligonucleotides. The drawbacks of these fluorescence in-situ hybridization (FISH)-based approaches, however, are the need for cell fixation and permeabilization, which can relocalize or extract cellular components(Fox et al., 1985; Schnell et al., 2012), the difficulty of assigning RNAs to specific organelles or cellular landmarks due to spatial resolution limits, and the limited information content compared to RNA sequencing: FISH and MERFISH can identify gene products, but further information, such as RNA isoform, UTR sequence, and mutations are lost. Finally, these transcriptome-wide imaging methods are challenging to perform and require specialized instrumentation not available to most laboratories. A related technique, *in situ* sequencing(Lee et al., 2014; Wang et al., 2018), also shows great promise, but is even more technically out of reach to most laboratories.

A recent adaptation of ribosome profiling(Ingolia et al., 2009) has enabled this technique to profile actively-translated mRNAs in specific cellular locales, namely the ER membrane in yeast and mammalian cells(Jan et al., 2014), and the mitochondrial outer membrane in yeast(Williams et al., 2014). While the spatial specificity of this live-cell approach is high, the methodology cannot, by its design, detect non-coding RNAs or silent (non-translated) mRNAs. Proximity-specific ribosome profiling is also not yet a fully generalizable method, as the requirement for biotin starvation during cell culture is prohibitively toxic to many organelles and cell types (such as mitochondria in mammalian cells, which has not been successfully studied by this approach – J. Weissman, personal communication).

Hence, there remains a need for new methodology that can map the spatial localization of thousands of RNAs at once, in living cells, and with minimal toxicity. The method should be applicable to any subcellular region, and capture full sequence details of any RNA type, enabling comparisons across RNA variants and isoforms. Here we develop the “APEX-seq” methodology in an effort to provide these capabilities. We develop and characterize the APEX-seq approach, and then apply it to nine distinct subcellular locations, generating highly specific maps of endogenous RNA localization in living human HEK (human embryonic kidney) 293T cells. Our data reveal correlations between RNA localization and corresponding protein localization, as well as RNA localization and underlying genome architecture. An analysis of mRNAs at the outer mitochondrial membrane suggests distinct mechanisms for RNA targeting that correlate with the sequence and function of the encoded mitochondrial proteins. These examples illustrate the versatility of APEX-seq and its ability to nominate and/or test novel biological hypotheses.

## RESULTS

### Development of APEX-seq Methodology

To meet the challenge of a generalizable method for RNA mapping in live cells, we considered a method that we previously developed for proteomic mapping(Lam et al., 2014; Rhee et al., 2013). APEX2(Lam et al., 2014) is a mutant of soybean ascorbate peroxidase that catalyzes one-electron oxidation of the membrane-permeable small molecule substrate biotin-phenol (BP). The resulting BP radical is very short-lived (half-life < 1 msec(Mortensen and Skibsted, 1997; Wishart and Rao, 2010)) and covalently attaches onto protein sidechains, enabling APEX2 to catalyze the promiscuous biotin-tagging of endogenous proteins within a few nanometers of its active site in living cells. Subsequently, biotinylated proteins are enriched with streptavidin and identified by mass spectrometry. The high spatial specificity of this approach has enabled APEX to be used not only for mapping organelle proteomes(Han et al., 2017; Hung et al., 2017; Hung et al., 2016; Hung et al., 2014; Lam et al., 2014; Loh et al., 2016; Rhee et al., 2013) but also protein interaction networks in living cells(Bersuker et al., 2018; Gupta et al., 2018; Lobingier et al., 2017; Markmiller et al., 2018; Paek et al., 2017).

We previously explored the extension of APEX to RNA by performing live cell proteome biotinylation, and then crosslinking endogenous RNAs to biotinylated proteins using formaldehyde (“APEX-RIP”)(Kaewsapsak et al., 2017). This enabled subsets of RNAs to be enriched using streptavidin and identified by RNA-seq(Mortazavi et al., 2008). We found that while APEX-RIP worked well in membrane-enclosed organelles such as the mitochondrial matrix, the spatial specificity was very poor in “open”, non-membrane enclosed cellular regions. For instance, RNAs enriched by APEX-ERM (APEX targeted to the ER membrane facing cytosol) were no different from those enriched by cytosolic APEX(Kaewsapsak et al., 2017). Two factors likely contribute to poor spatial specificity: formaldehyde crosslinking across many molecules, and the long 16-minute labeling protocol which gives ample time for APEX-biotinylated proteins to redistribute and crosslink to distal RNAs.

A more straightforward and potentially higher specificity approach would be to bypass formaldehyde crosslinking altogether and use APEX peroxidase to directly biotinylate cellular RNAs within a short time window (Figure 1A). Previous studies have shown that phenoxyl radicals can react with electron-rich nucleobases such as guanine; this chemistry forms the basis of mutagenicity of phenolic xenobiotics(Dai et al., 2005; Dai et al., 2003a, b), but it has not previously been explored for live-cell RNA labeling. To test if peroxidase-generated phenoxyl radicals could biotinylate RNA in vitro, we combined horseradish peroxidase (HRP), which catalyzes the same one-electron oxidation reaction as APEX2(Loh et al., 2016), with tRNA, biotin-phenol (BP), and H_2_O_2_. On a streptavidin dot blot, we observed robust tRNA biotinylation that was abolished by RNase treatment, but unaffected by proteinase K treatment, which degrades proteins (Figure S1A). We then asked whether covalent BP modification of RNA would block the progress of a reverse transcriptase (RT). 5S ribosomal RNA labeled with HRP, BP, and H_2_O_2_ was extended using a ^32^P-labeled primer. Figure S1D and E show that while full-length transcripts are still produced, multiple RT stops are observed at G-rich regions in peroxidase-labeled RNA samples, but not in a negative control. We performed additional experiments to characterize the covalent adduct formed with G in vitro by HPLC and mass spectrometry (Figure S1B and C).

**Figure 1:**
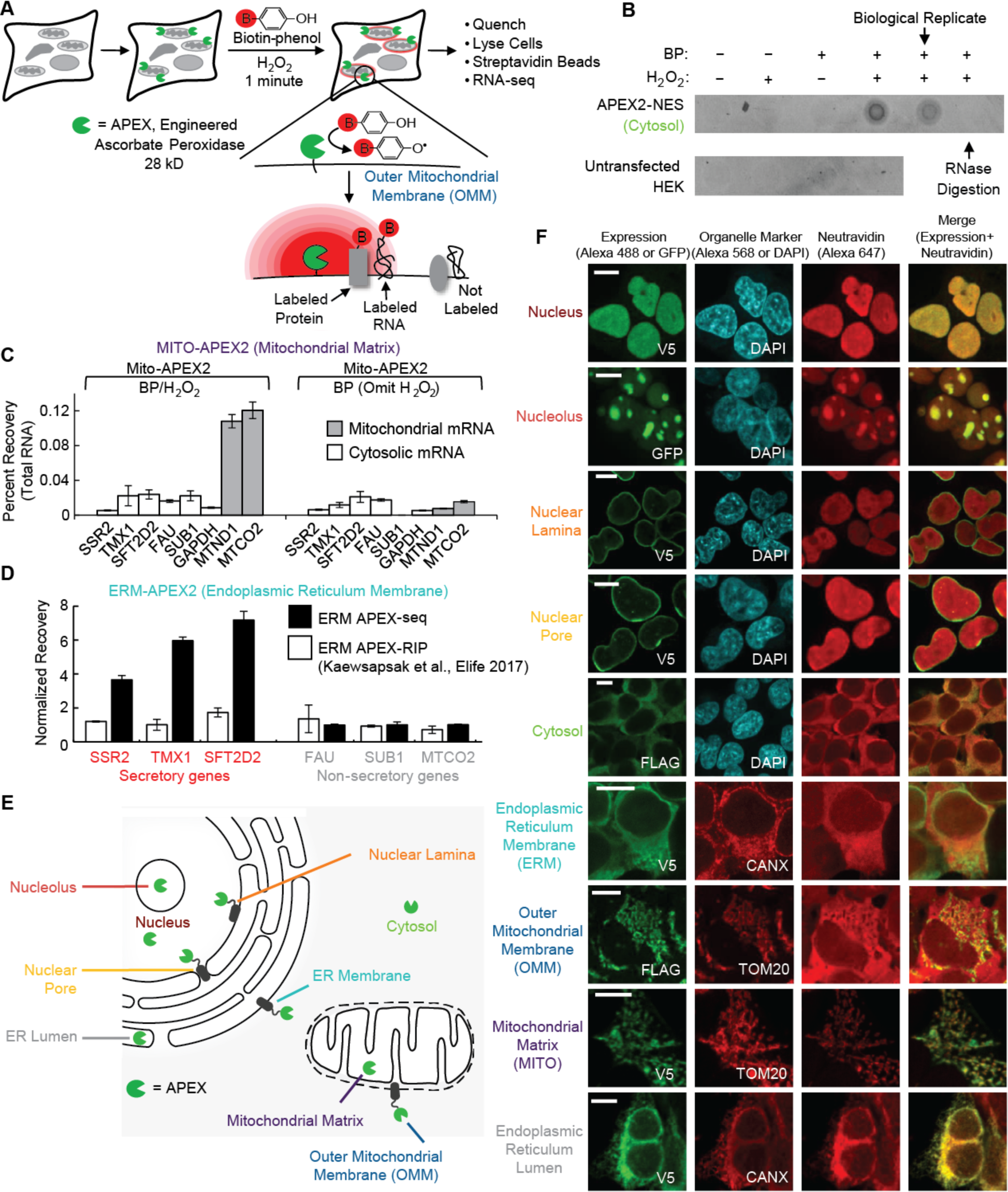
Development of APEX-seq methodology. (A) APEX2-mediated proximity biotinylation of endogenous RNAs. APEX2 peroxidase(Lam et al., 2014) is genetically targeted to the cellular region of interest. Addition of biotin-phenol (red B = biotin) and H_2_O_2_ to live cells for 1 minute results in covalent biotinylation of endogenous proteins and RNA within a few nanometers of APEX2. APEX-generated biotin-phenoxyl radicals have a half-life of < 1 millisecond. After 1 minute labeling in live cells, the reaction is quenched, and cells are lysed. Biotinylated RNAs are separated using streptavidin-coated beads, polyA-selected, and analyzed by RNA-seq. (B) Streptavidin-biotin dot blot analysis of direct RNA biotinylation by APEX2 in cells. HEK-293T cells stably expressing APEX2 in the cytosol (FLAG-APEX2-NES) were labeled with BP and H_2_O_2_ for 1 minute, then the RNA was extracted and blotted (500 ng total RNA per condition). Only when BP, H_2_O_2_, and APEX2 were all present was streptavidin signal observed. RNase treatment of the sample prior to dot blot abolished the signal. Figure S1A-E present more evidence of direct biotin labeling of RNA during the peroxidase-catalyzed reaction. (C) RT-qPCR analysis showing specific enrichment of mitochondrial RNAs (grey) over cytosolic mRNAs (white) using mito-V5-APEX2. HEK-293T cells stably expressing APEX2 targeted to the mitochondrial matrix were labeled for 1 minute with BP and H_2_O_2_. Biotinylated RNAs were enriched with streptavidin beads following total RNA extraction and then analyzed by RT-qPCR. Data are the mean of four replicates-± one standard deviation. Figure S1F shows the optimization of APEX-seq. (D) RT-qPCR analysis showing specific enrichment of secretory (red) over non-secretory (grey) mRNAs with APEX-seq, but not APEX-RIP. HEK-293T cells stably expressing APEX2 targeted to the ER membrane (facing cytosol) were labeled for 1 minute with BP and H_2_O_2_. For APEX-seq, biotinylated RNAs were enriched with streptavidin beads following total RNA extraction and then analyzed by RT-qPCR. For APEX-RIP, RNAs were crosslinked to proteins for 10 minutes before streptavidin beads enrichment. Data are the mean of four replicates-± one standard deviation. The data was normalized such that the mean enrichment of non-secretory RNAs was 1 for both APEX-seq and APEX-RIP. (E) Human cell showing nine different subcellular locations investigated by APEX-seq. ER, endoplasmic reticulum. (F) Fluorescence imaging of APEX2 localization and biotinylation activity. Live-cell biotinylation was performed for 1 minute with BP and H_2_O_2_ in HEK-293T cells stably expressing the indicated APEX2 fusion protein. APEX2 expression was visualized by GFP or anti-V5/FLAG staining (green). Biotinylated species were visualized by staining with neutravidin-AlexaFluor 647 (red). DAPI is a nuclear marker. Antibodies against endogenous *TOM20* and *CANX* were used as markers for the mitochondria and ER, respectively. The nuclear-pore construct had the APEX2 moiety facing the nucleus, as judged by biotin labeling. Scale bars, 10 µm.

To test APEX-catalyzed RNA biotinylation in living cells, which contain millimolar concentrations of the BP radical quencher glutathione that could impair labeling, we generated HEK-293T cells stably expressing APEX2 in the cytosol. We labeled the cells with BP and H_2_O_2_ for 1 minute, extracted total RNA (leaving behind protein), and analyzed the RNA by streptavidin dot blot. Figure 1B shows that biotin signal corresponding to labeled RNA disappears upon omission of BP or H_2_O_2_, or treatment with RNase. This result, combined with the RT-stop experiment above, suggests that APEX directly tags RNA with biotin, not merely biotinylating RNA-associated proteins. Separately, we confirmed that RNAs remain intact after APEX labeling (Figure S1G).

APEX-catalyzed RNA biotinylation in cells proved to be highly specific. We first used HEK cells expressing APEX2 in the mitochondrial matrix (MITO-APEX2), because the mitochondrial transcriptome is well-understood via sequencing of the mitochondrial genome (mtDNA). After 1 minute of BP labeling, cells were lysed, and biotinylated RNAs enriched with streptavidin beads. We optimized a series of denaturing washes to remove all species (RNA or protein) that were not *themselves* directly biotinylated (Figure S1F). Enriched material was then eluted via digestion of streptavidin with proteinase K, and specific RNAs were detected by RT-qPCR. Figure 1C shows enrichment of mtDNA-encoded mRNAs (*MTND1* and *MTCO2*), but not negative-control cytosolic mRNAs that are encoded by the nuclear genome (e.g., *GAPDH, SSR2, TMX1*).

However, because the mitochondrial matrix is enclosed by a tight membrane that is impervious to BP radicals(Rhee et al., 2013), it does not provide a rigorous test of APEX-seq labeling radius. To evaluate the spatial specificity of labeling, we examined the ER membrane (ERM), which we previously attempted but failed to map using APEX2-ERM and APEX-RIP(Kaewsapsak et al., 2017). When we performed a comparison of APEX-seq and APEX-RIP using HEK 239T cells expressing APEX on the ER membrane (facing the cytosol), we found that APEX-seq could clearly enrich secretory mRNAs on the ER membrane over cytosolic mRNAs, whereas APEX-RIP could not (Figure 1D). Given the very close proximity between ER membrane-associated RNAs and cytosolic RNAs that are immediately adjacent to the ER, these results suggest that APEX-seq has a labeling radius of only a few nanometers.

### Validation of APEX-seq

Encouraged by the results above, we expanded APEX-seq to nine distinct subcellular locations (Figure 1E). Correct localization of each APEX2 fusion protein (domain structures shown in Figure S2A) in HEK-293T cells was confirmed by imaging with organelle markers (Figure 1F). We prepared two replicates for each location (Figure S2B and S2C, correlation coefficient *r* between replicates ranged from 0.97 to 1) as well as negative controls in which H_2_O_2_ was omitted from the labeling reaction. After labeling for 1 minute and cell lysis, we enriched biotinylated RNAs and prepared APEX-seq libraries with polyA+ selection. Transcripts shorter than 100 nt were excluded during purification. We used DESeq2(Love et al., 2014) for data analysis and used a q-value (FDR-adjusted p-value) < 0.05 and a log_2_ fold-change > 0.75 to select for significantly enriched transcripts.

Consistent with the RT-qPCR results for the mitochondrial matrix shown in Figure 1C, our sequencing-based analysis showed that all 13 mRNAs and 2 rRNAs encoded by mtDNA are strongly enriched by MITO-APEX2 labeling (Figure 2A, S2D and S2E, mean enrichment > 11-fold), whereas no mRNAs encoded by the nuclear genome are enriched. Figure 2B shows good correlation (*r* > 0.9) between technical replicates.

**Figure 2:**
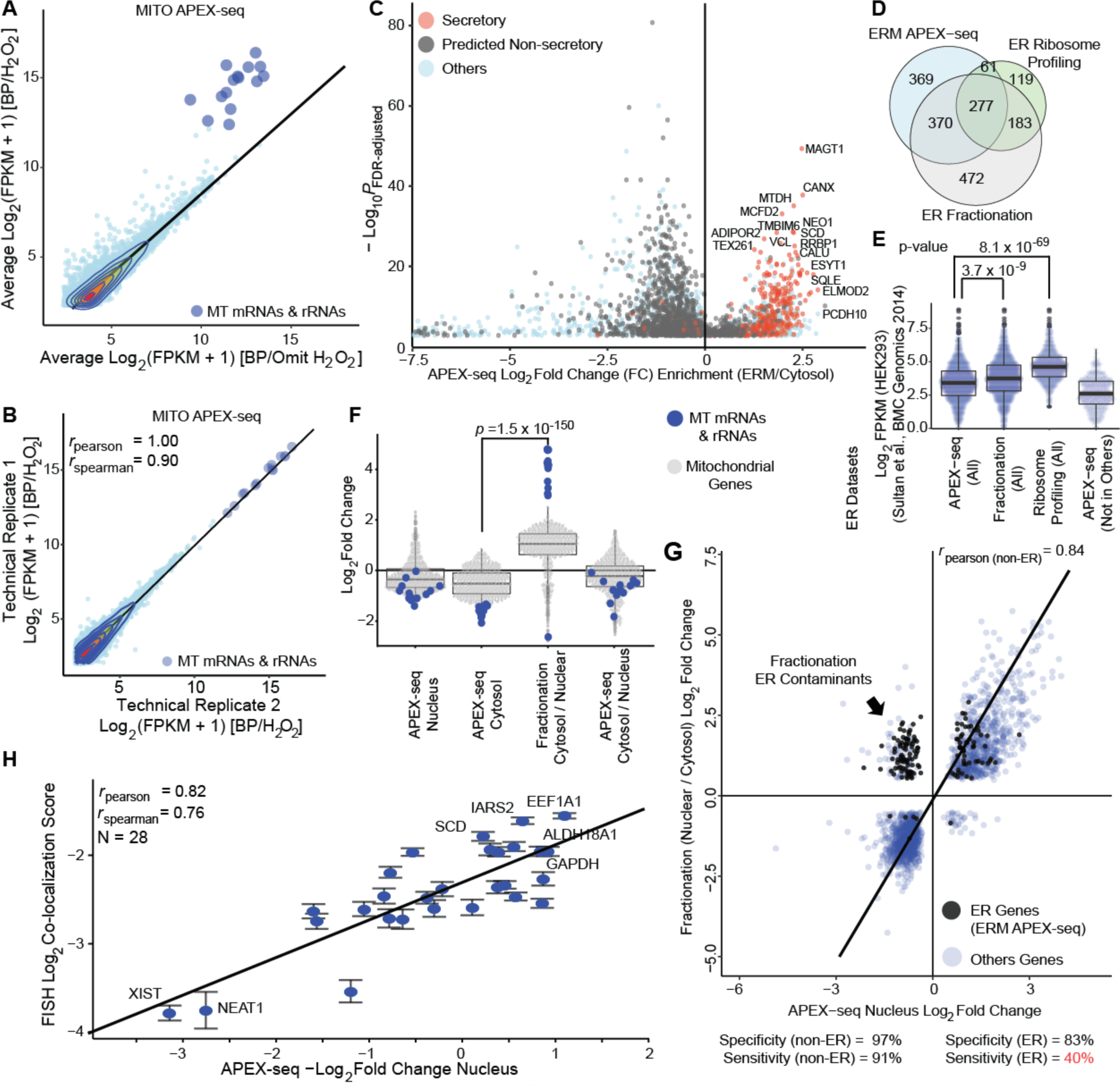
Validation of APEX-seq. (A) APEX-seq in the mitochondrial matrix. Transcript abundance in experiment (y axis) plotted against negative control (x axis, omit H_2_O_2_). All 13 mRNAs and 2 rRNAs encoded in the mitochondrial genome (large blue dots) are enriched by APEX (mean enrichment > 11-fold). FPKM, fragments per kilobase of transcript per million reads. More than half of the reads map to these mRNAs in the experiment compared to ~5% in the controls. Figure S2A shows the individual constructs generated to specifically target APEX2 to the mitochondria, as well other locations. Figure S2B-C shows the APEX-seq mapping statistics and good agreement between biological replicates. Figure S2D-G present more details on enrichment of mitochondrial rRNAs and mRNAs by APEX-seq. (B) Scatter plot of transcript abundance in the mitochondrial matrix (MITO) APEX-seq shows good agreement between technical replicates. (C) APEX-seq at the ER membrane (ERM), facing cytosol. Volcano plot showing APEX-catalyzed enrichment (positive fold-change) of secretory mRNAs (red, defined as mRNAs previously enriched by proximity-specific ribosome profiling at the ER membrane(Jan et al., 2014)) over non-secretory mRNAs (black, defined as non-secretory by Phobius, SignalP, and TMHMM. Figure S2H-K provide further details on the specificity and coverage of APEX-seq ERM dataset. (D) Comparison of ERM-enriched RNAs by APEX-seq, proximity-specific ribosome profiling(Jan et al., 2014), and ER fractionation-seq(Reid and Nicchitta, 2012). Almost 2/3^rd^ of the RNAs recovered by APEX-seq were also obtained by the other two methods. (E) Transcript abundance (FPKM) analysis of genes enriched by ERM APEX-seq, fractionation-seq, proximity-specific ribosome profiling, and genes unique to the APEX-seq dataset. FPKM data in HEK 293T cells were taken from a published dataset(Sultan et al., 2014). (F) APEX-seq in the cytosol does not recover intra-mitochondrial RNAs whereas nuclear fractionation-seq does. The 13 mRNAs and 2 rRNAs encoded by the mitochondrial genome are shown as blue dots. The cytosolic fraction of bulk fractionation RNA-seq is enriched in these mitochondrial contaminants, whereas cytosolic and nuclear APEX-seq datasets are not. P-value is from a Mann-Whitney U test. Figure S2B shows the sequence-mapping statistics and agreement between technical and biological replicates. (G) APEX-seq yields cleaner results than bulk fractionation RNA-seq. Nucleus APEX-seq fold changes are highly correlated with bulk fractionation RNA-seq when considering non-ER genes (blue points, obtained by excluding ERM APEX-seq enriched genes). However, bulk fractionation suffers from contamination by ER transcripts (black points) that are depleted in APEX Seq, which results in lower agreement between the two methods for ER transcripts. Using fractionation RNA-seq data as training set, we could compute an estimate of accuracy and precision by nucleus APEX-seq. For non-ER transcripts, APEX Seq yields both high precision and accuracy. Figure S2L-M show further comparisons of APEX-seq versus fractionation data. (H) Correlation between nucleus APEX-seq enrichment and FISH imaging colocalization score for 28 selected genes. FISH co-localization score was calculated as the percentage of total signal overlapping with a mitochondrial marker, *MTND3*. Figure S3A shows individual sequential FISH images. Figure S3B shows the correlation between FISH score and OMM/nucleus APEX-seq enrichment.

To assess APEX-seq in an “open” subcellular region (not enclosed by a membrane), we focused again on the ER membrane (ERM). The ERM is particularly valuable for methodology validation because there are well-established “true positives” (mRNAs encoding secreted proteins that are known to be translated at the ERM) and well-established “false positives” (mRNAs encoding soluble cytosolic proteins). In addition, the ERM-associated transcriptome has been mapped by previous methods, including fractionation-seq(Reid and Nicchitta, 2012) and proximity-specific ribosome profiling(Jan et al., 2014), allowing a head-to-head comparison with APEX-seq. Our RT-qPCR experiment above using APEX2-ERM (Figure 1D) was encouraging, but we wished to perform a more comprehensive analysis of APEX-seq via unbiased sequencing of the ERM-associated transcriptome.

From reliable sequencing data for over 17000 Ensembl genes(Hubbard et al., 2002), we observed that ERM APEX-seq strongly enriched mRNAs encoding secretory proteins over non-secretory mRNAs (*p*<10^−100^, Figure 2C). Using ROC analysis as previously described(Linden, 2006) and shown in Figure S2H, we determined quantitative cut-offs, and arrived at an ERM APEX-seq dataset of 1077 RNAs. To estimate the specificity of this dataset, we calculated the fraction of genes with prior secretory annotation in Gene ontology cellular component (GOCC), SignalP(Petersen et al., 2011), TMHMM(Krogh et al., 2001) or Phobius(Kall et al., 2004)). 90% of our ERM APEX-seq mRNA genes code for known secretory proteins; this is therefore an estimate of the lower bound for the specificity of our dataset. The remaining 10% of genes (107 mRNA genes) could be newly discovered ERM-associated RNAs (identified as such by ERM APEX-seq but not by previous methods), or they could be false positives.

We assessed sensitivity, or depth-of-coverage, for our ERM APEX-seq dataset by calculating the fraction detected for a hand-curated list of 71 very well-established ER-resident proteins. APEX-seq recovered 73% of these genes. To enable comparison, we analyzed previous fractionation-seq(Reid and Nicchitta, 2012) and ribosome profiling(Jan et al., 2014) ER datasets. We found that specificity and coverage (Figure S2J) of each were comparable to that of APEX-seq, but the Venn diagram (Figure 2D) shows that each method recovers somewhat different subsets of transcripts. Further analysis of genes enriched by APEX-seq but *not* enriched by fractionation-seq or ribosome profiling show that many of these are lower in total cellular abundance (Figure 2E). Hence, the three methods appear to be complementary, at least for the ERM, and APEX-seq may be better able to recover lower abundance transcripts.

While the majority (97%) of ERM APEX-seq-enriched species are mRNAs, this dataset highlights a few noncoding RNAs - including antisense-RNAs (CKMT2-AS1, RAMP2-AS1 and PINK1-AS) and lncRNAs (TUG1 and NORAD) which are not detected by either biochemical fractionation or ribosome profiling. These RNAs remain invisible to ribosome profiling as they are not translated, and because they also tend to be lowly expressed (Figure 2E) they may be easily missed by fractionation. Our results thus highlight some advantages offered by APEX-seq.

Apart from the ERM, we also used the nuclear and cytosolic compartments to benchmark APEX-seq. For comparison, we used the same HEK 293T cells to prepare our own biochemically-purified nuclear and cytosolic fractions (following protocols published in Gagnon et al., 2014), and then analyzed them by RNA-seq in the same manner as the APEX-seq samples (to produce our own “fractionation-seq” data). We observed that our nuclear and cytosolic APEX-seq data were much more specific than our corresponding fractionation-seq data. For instance, our APEX-seq gene lists lacked the mitochondrial matrix contaminants prevalent in the cytosolic fractionation data (Figure 2F) as well as the ER contaminants prevalent in the nuclear fractionation data (Figure 2G and Figure S2M).

Excluding the known ER contaminants in the nuclear fractionation-seq dataset, we compared the remaining genes to APEX-seq in order to estimate the accuracy and precision of our methodology. We found that nuclear APEX-seq fold-changes were highly correlated with fractionation-seq nuclear enrichment (r_pearson_ = 0.84). Furthermore, relative to fractionation-seq, APEX-seq was accurate (94%), precise (96%), specific (97%) and sensitive (91%). The corresponding values for all genes (including ER contaminants in the nuclear fractionation-seq) were 90%, 96%. 97% and 83% respectively. We also observed that the RNA length distributions in nuclear fractionation and APEX-seq are very similar (Figure S2L), suggesting an absence of length bias for APEX-seq.

For final validation of APEX-seq, we selected 29 RNAs that show differential localization to cytosol versus nucleus and analyzed them by FISH imaging(Chen et al., 2015b) (Figure S3). Figure 2H shows strong correlation (*r* = 0.82) between nuclear APEX-seq depletion and FISH colocalization with a mitochondrial marker MTND3 in the cytoplasm, as expected. In addition, transcripts with high OMM (outer mitochondrial membrane) APEX-seq enrichment display higher colocalization with MTND3 as well (Figure S3B).

Together, the comparisons of mitochondrial matrix, ERM, nuclear, and cytosolic APEX-seq data to published databases and studies, as well as to our own fractionation-seq and FISH imaging data, suggest that APEX-seq provides accurate (specific and sensitive) RNA localization in multiple subcellular compartments.

### APEX-seq Reveals Transcriptome-wide Spatial Organization

APEX-seq from nine compartments produced a subcellular map of over 3200 RNAs and revealed their detailed spatial organization (Figure 3A). RNAs broadly partitioned into four general categories of localization: (1) nuclear (encompassing nucleolus, nucleus and nuclear lamina), (2) mitochondrial membrane/ER, (3) cytosol, and (4) the remaining (which includes ER lumen, mitochondrial matrix, and nuclear pore) (Figure 3A). Each transcript further localized to just one or two locations within each general category (Figure 3B, 3G, Methods). For each cluster of co-localized transcripts, Gene Ontology (GO) analysis identified enriched cellular components that are consistent with the annotated locations (Figure S3D). For example, clusters 1-2 that include ERM- and OMM-enriched RNAs are enriched for secretory and membrane-associated genes, whereas clusters 3-4 that largely include nuclear RNAs are enriched for nuclear-associated GO-terms.

**Figure 3:**
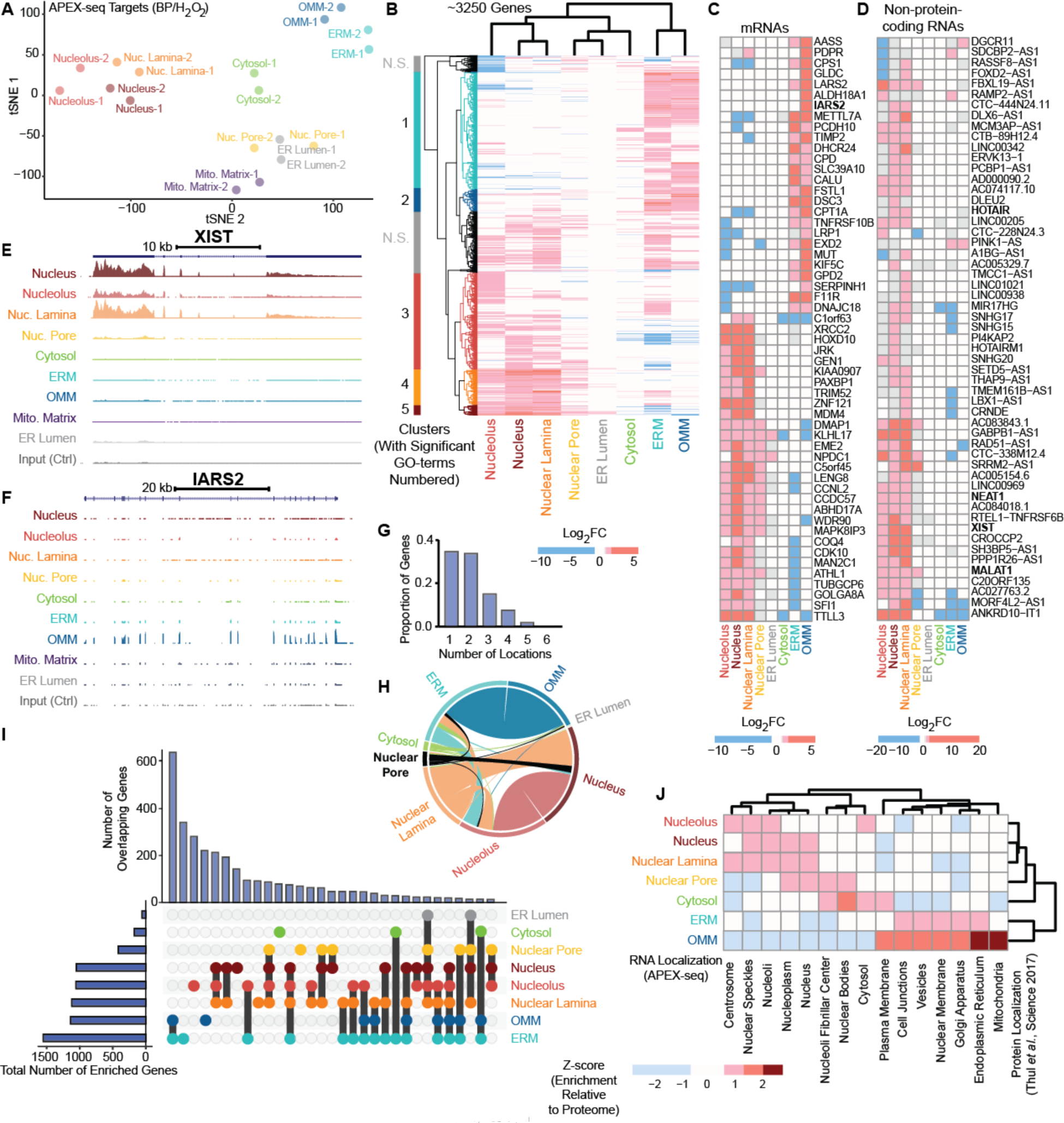
Analysis of subcellular transcriptome maps. (A) T-distributed Stochastic Neighbor Embedding (t-SNE) plot showing the separation and clustering of APEX-seq libraries. Within the labeled samples (with BP/ H_2_O_2_), the nuclear locations (nucleus, lamina, nucleolus) cluster together; and the ER and OMM form a cluster. The nuclear pore clusters separately from both the nuclear and cytoplasmic components. (B) Heatmap of transcripts enriched by APEX-seq showing clustering of the 3262 genes that specifically localize to at least one location and have fold-change data from all locations. The nuclear locations (nucleolus, nucleus and nuclear lamina) cluster together, and separate from the ERM and OMM. The nuclear-pore transcriptome is closer to the cytosol, and strikingly different from the nuclear lamina. Figure S3D shows the GO-terms associated with the clusters identified. Figure S4 shows the overlap of ERM and OMM. (C) Heatmap showing the APEX-seq fold changes for the mRNA transcripts found to be most variable among the locations investigated. Many of these RNAs localize to the ERM and OMM, as well as to nuclear locations. (D) Heatmap showing the APEX-seq fold changes for non-coding RNAs (excluding pseudogenes) that have the most-variable localization enrichment. Almost all these RNAs are localized within the nucleus. A few well-known noncoding RNAs (*XIST*, *MALAT1*, *HOTAIR* and *NEAT1*) are shown in bold. (E) (F) Genome tracks for (E) *XIST*, a nuclear non-coding RNA, and (F) *IARS2*, an mRNA encoding a mitochondrial tRNA synthetase, across 8 different subcellular locations. For each location, the reads were averaged across two APEX-seq replicates (after normalization of each to the same read depth of 35 million). The control tracks at bottom were generated by averaging 18 controls from all 9 constructs (2 replicates per location). Consistent with previous studies, *XIST* preferentially localizes to the nuclear lamina over the nucleolus or nuclear pore, and *IARS2* localizes to the OMM. (G) Of the ~3250 genes analyzed, most localize to only 1 or 2 of the 8 locations (excluding mitochondrial matrix) interrogated. (H) Circos plot showing the co-localization of RNAs to multiple locations. We examined RNAs that are significantly enriched in at least one location and have fold-change data from all locations. (I) Transcripts overlapping in multiple locations, as determined by APEX-seq. We observe large overlap between the ERM- and OMM-localized transcripts, and between the nuclear locations (nuclear lamina, nucleolus, and nucleus). Of these ~3250 RNAs that show significant local enrichment, the ERM has the most RNAs, while the ER lumen has the fewest. As we normalize our data relative to the unlabeled transcriptome i.e. effectively whole-cell (Figure S6F), we recover comparatively few RNAs (~173) by cytosol APEX-seq when we use the same log_2f_ old-change cutoff (0.75) used for other challenging locations. Because the cytosol RNAs constitute a majority of the RNAs in the cell (as confirmed by cytosol/nuclear fractionation), it is difficult for RNAs to show a significant (log_2f_old-change > 0.75, *p*_adjusted_ < 0.05) enrichment in the cytosol (Figure S3C). Nonetheless the RNAs that show any enrichment by cytosol APEX-seq are significantly enriched in the cytosol fractionation-seq data relative to transcripts depleted by cytosol APEX-seq (*p* < 10^−100^). Use of an alternative RNA reference pool, such as the nuclear APEX-seq, can readily highlight cytoplasmic RNAs. (J) Heatmap showing the protein localization of the transcripts enriched by APEX-seq. Here the scale shows enrichment of protein categories relative to entire proteome database(Thul et al., 2017). We find that clustering based on the protein recapitulates the similarity in ERM and OMM, relative to the nuclear components. The data suggests that there is a correlation between protein and RNA localization. Figure S4A shows this information in more detail.

We find many mRNAs that are primarily localized to one of the cytosolic locations or one of the nuclear locations. Some cytosolic examples include mRNAs encoding mitochondrial proteins(Calvo et al., 2015) (*AASS, LARS2*) as well as known ER proteins(Jan et al., 2014; Reid and Nicchitta, 2012) (*CALU, CPD*). The nuclear mRNA transcripts include known nuclear-enriched RNAs(Sultan et al., 2014) (*WDR90, TTLL3, NPDC1*) (Figure 3C). In contrast to mRNAs, lncRNAs (long noncoding RNAs) are predominantly nuclear; these include extensively-studied lncRNAs such as HOTAIR, NEAT1, XIST and MALAT1 (Figure 3D), which have also been mapped in previous studies(Cabili et al., 2015).

Inspection of APEX-seq data of individual transcripts on the UCSC genome browser (Kent et al., 2002) provides potential insights into their function. For example, APEX-seq data showed that XIST (X-inactive specific transcript), a nuclear lncRNA, is enriched at the nuclear lamina but not the nearby nuclear pore (Figure 3E). These findings are consistent with the known role of XIST in coating the inactive X chromosome in female cells(Penny et al., 1996), leading to transcriptional silencing and localization of the inactive X to the nuclear lamina(Chen et al., 2016). Another example is *IARS2* (mitochondrial isoleucyl tRNA synthetase 2 that is encoded by the nuclear (not mitochondrial) genome), whose mRNA was enriched by APEX-seq at the OMM, but not ERM or mitochondrial matrix (Figure 3F). Because *IARS2*’s protein product is known to reside in the mitochondrial matrix, the APEX-seq data suggest a staged production process in which protein translated at one location (e.g., OMM) is imported into another (e.g., mitochondrial matrix), a point that we further explore in Figure 7.

We used our APEX-seq map to explore the relationship between mRNA localization and the location of the protein product at the steady state. Comparison to the Protein Cell Atlas database(Thul et al., 2017) revealed remarkable concordance between RNA and protein localization (Figure 3J, Figure S4A). For example, the ER transcriptome preferentially codes for proteins that localize to the ER, golgi and vesicles, rather than proteins that localize to the nucleus, nucleolus or cytosol. Less expectedly, our data also show that mRNAs enriched in nuclear locations tend to code for proteins enriched in nuclear speckles, nucleoplasm, but not the plasma membrane (Figure 3J). Although for the nuclear compartments the correlation between RNA and protein localization is not as strong as for the ERM and OMM, the correlation is nonetheless surprising if protein translation occurs exclusively in the cytosol. A previous study using fractionation and imaging found that many spliced, polyadenylated mRNA (i.e. mature) transcripts are highly abundant in the nucleus(Bahar Halpern et al., 2015), and the authors show that this mRNA nuclear retention buffers cytoplasmic transcript levels from noise that emanates from transcriptional bursts. Thus, mRNAs in the nucleus might serve as “reserve pools” or “holding stations” that help to dampen gene-expression noise. We build on these findings by speculating that nuclear-destined proteins, which are highly-enriched for nucleic-acid binding proteins (FDR < 5 × 10^−13^, GO-term biological process) whose concentrations may have to be precisely tuned, may have mRNAs that are retained in nuclear subcompartments in order to better shield the amount of mRNA available for translation from noise.

In each of the following sections, we use our APEX-seq resource to explore and/or develop a number of biological hypotheses.

### Nuclear Pore As a Staging Area for RNA Export

RNA transcripts must pass through the nuclear pore to go from their production sites in the nucleus into the cytoplasm. Previous studies have suggested that the nuclear pore may act as a staging area for cytoplasm-destined transcripts(Wickramasinghe and Laskey, 2015). For example, the nuclear pore subunit protein TAP/p15 has been shown to interact with several exonjunction-complex (EJC) components that are involved in mRNA splicing(Katahira, 2015; Singh et al., 2012). Thus there exists an intimate connection between export and splicing(Reed and Cheng, 2005).

Our nuclear pore APEX-seq construct targets APEX2 to SENP2 (Sentrin-specific protease 2), which is known to interact with NUP13, a component of the nuclear-pore basket(Walther et al., 2001). Our APEX-seq data reveal a remarkable resemblance between the set of transcripts at the nuclear face of the nuclear pore and RNAs in the cytoplasm (Figure 3B), in contrast to RNAs from other locations within the nucleus (Figure 3A). This is a striking finding because the nuclear pore is physically embedded in the nuclear lamina, and their APEX-mediated biotinylation patterns appear similar by immunofluorescence (Figure 1F). Yet APEX-seq identifies striking differences in transcripts localized to the nuclear pore versus the nuclear lamina, showing that RNA content can vary within the cell just nanometers apart.

Because transcripts enriched at the nuclear-pore resemble those in the cytosol (Figure 3B), these findings suggest that the inward facing domain of the nuclear pore is a staging area where only properly spliced and sorted transcripts ready to export to cytoplasm are allowed to congregate. In other words, the dwell time of properly spliced and processed RNAs at the nuclear pore is systematically much larger than incompletely-processed RNAs, thereby allowing us to enrich the former class by APEX-seq. The nuclear pore may be akin to an airport terminal, where only passengers with proper tickets and passports are allowed to board. Blobel famously conceived of the nuclear pore as a “gene gate”(Blobel, 1985), where all transcripts from a given gene locus would depart from a specific nuclear pore and be delivered to a pre-defined cytoplasmic location(Fasken and Corbett, 2009; Kim et al., 2018). More recent studies have refined the “gene gating” concept(Brown and Silver, 2007), with mRNA transcripts associating with a number of proteins to form a messenger ribonucleoprotein (mRNP) particle as they are shuttled from the nucleus to the cytosol. Our results support the prevailing consensus that the nucleoplasmic milieu of the nuclear pore complex (NPC) has a critical role in mRNA surveillance, which prevents partially spliced polyadenylated transcripts (Figure 4A-C) from reaching the cytoplasm.

**Figure 4:**
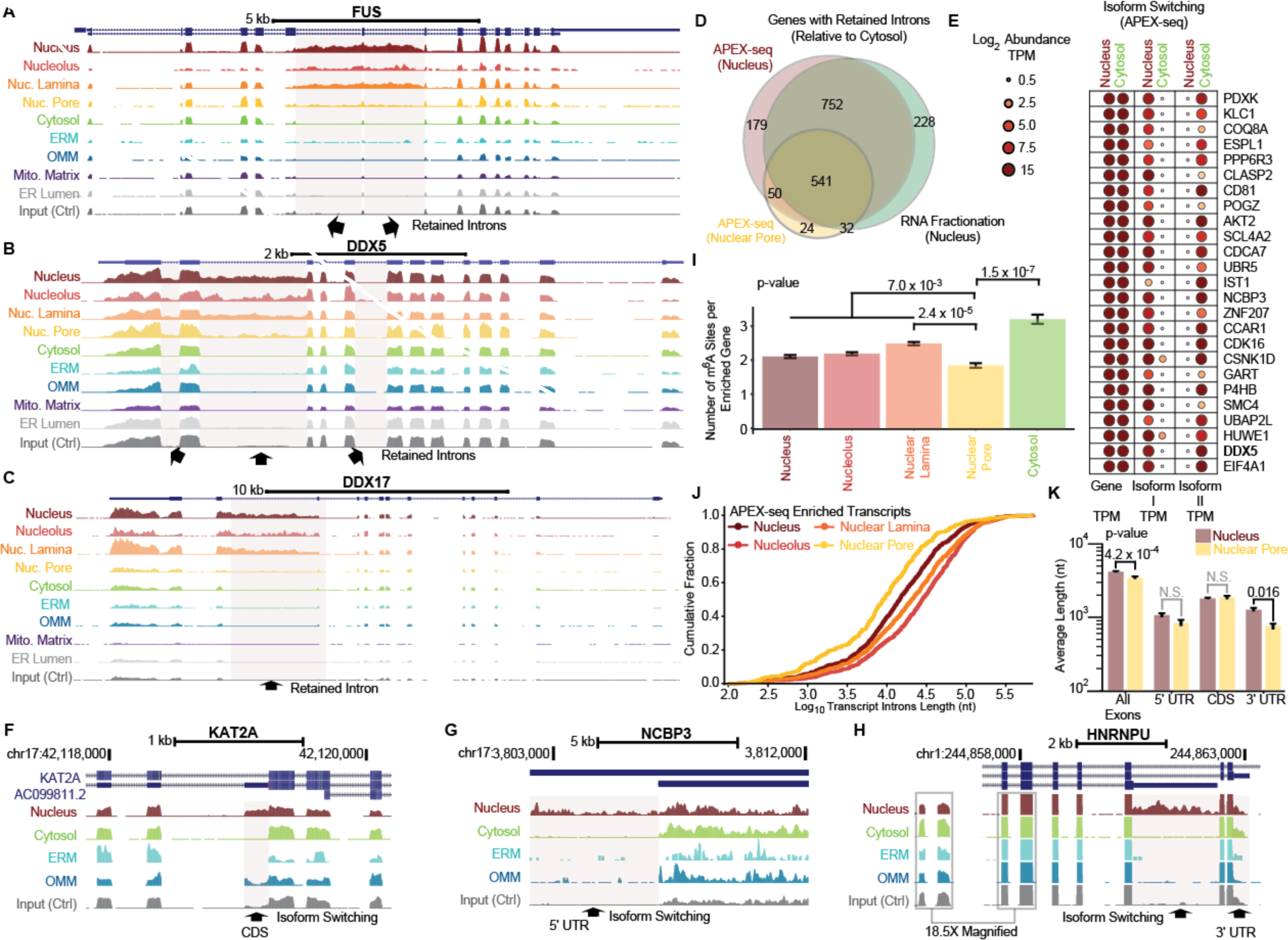
APEX-seq reveals principles related to RNA isoforms and introns. (A) (B)(C) Unlike MERFISH and other imaging-based methods, APEX-seq can identify splice isoforms and retained introns. The genome tracks of (A) *FUS*, an mRNA encoding a protein involved in aggregation and ALS, reveal transcripts with retained introns in the nuclear sub-compartments that do not make it to the nuclear pore and beyond. (B) and (C) show the genome tracks of two other transcripts, *DDX5* and *DDX17*, with retained introns. (D) Fractionation-seq (green) and nucleus APEX-seq (red) identify roughly the same genes with retained introns. The nuclear-pore APEX-seq transcriptome has fewer retained introns relative to the nucleus. (E) Using APEX-seq, we can identify transcripts that are highly abundant in both cytosol and nucleus at the gene level, but switch isoforms at the transcript level. TPM, transcript per million. Figure S4A-E provide more details on how these transcripts were identified. (F) (G) (H) UCSC browser tracks showing examples of isoform switching across nuclear and cytosolic locations for (F) *KAT2A* (lysine histone acetyltransferase 2A) in a putative coding sequence (CDS), (G) *NCBP3* (nuclear cap-binding protein subunit 3) in the 5′ UTR and (H) *HNRNPU* (heterogenous nuclear ribonucleoprotein U) in the 3′ UTR respectively. (I) Number of m^6^ A present per transcript enriched by APEX-seq. High-confidence m6A sites in HEK293 were obtained from the literature(Meyer et al., 2012). P-values are from a Fisher’s exact test. (J) Cumulative distribution of the introns length for genes enriched by APEX-seq in the nuclear locations. We observe shorter transcripts at the nuclear pore relative to other locations. Here the transcript length was calculated by considering the most-abundance transcript isoform for each gene across all locations in the APEX-seq data. Figure S4J-L shows the same trends for the transcript exon lengths, as well as the corresponding distributions when calculating transcript length by considering the longest-stable isoform for each gene. (K) Barplots of average length of nuclear pore and nucleus enriched transcripts by mature transcript length (i.e. all exons), 5′ UTR, CDS (coding sequence) and 3′ UTR. P-values are from a one-sided Mann-Whitney U test. Errors are standard error of mean (S.E.M.).

### OMM and ERM Transcriptome Overlap

Another striking observation from our subcellular RNA map is the substantial overlap between the OMM and ERM transcriptomes (Figure 3H-I). Using a more-stringent ratiometric approach(Hung et al., 2014; Kaewsapsak et al., 2017), we confirmed that almost two-thirds of RNAs are shared by OMM and ERM, with almost 95% of shared messenger RNAs encoding secreted proteins (Figure S4B-C). This finding is not surprising given that the ER and mitochondria are often in close contact(Friedman et al., 2011; Giacomello and Pellegrini, 2016; Valm et al., 2017) and may be physically tethered to each other(Friedman et al., 2011; Giacomello and Pellegrini, 2016; Hung et al., 2017; Kornmann et al., 2009). Further analysis shows that relative to all secretory mRNAs, the cohort of OMM-proximal secretory mRNAs is not enriched for transcripts encoding transmembrane proteins (TMHMM analysis) but does appear to be strongly enriched for signal peptide (signalP)-encoding transcripts(Petersen et al., 2011) (Figure S4D-F). Actively translating ribosomes (polysomes) have previously been observed at mitochondria-rough ER contact sites(Williams et al., 2014). Our data raise the hypothesis that signal peptide-containing secreted proteins may be preferentially translated at these mito-rough ER junctions. Reassuringly, APEX-seq from the ER construct within the lumen recovered very few transcripts (Figure 3B, S3C).

### APEX-seq Reveals Differential Localization for Transcript Isoforms

The diversity of spatial localization occurs not only for RNAs encoding different genes, but also for transcript isoforms of the same gene (Figure 4A-D). For example, *FUS* (fused in sarcoma) mRNA, encoding a nuclear protein implicated in amyotrophic lateral sclerosis (ALS) and reversible phase separation(Patel et al., 2015), shows intron retention within the nuclear locations, but not the cytosolic ones (Figure 4A). Previous work has found such “retained introns” to often be evolutionarily conserved, stable (RNA half-life > 1 hr), not subject to nonsense-mediated decay (NMD), and enriched within mRNAs encoding RNA-binding proteins (RBPs)(Boutz et al., 2015). Figure 4B-C shows two additional mRNA genes, dead-BOX helicase 5 (*DDX5)* and dead-BOX helicase 17 (*DDX17)*, that contain retained introns. Importantly, *FUS* mRNA with retained intron is excluded from the cytosol by the nuclear pore, and APEX-seq of the nuclear pore has far fewer transcripts with retained introns compared to the nucleus genome-wide (Figure 4D). These results are again consistent with the role of the nuclear pore as a “gene gate” for RNA quality control. We note that the current model of EJC–nuclear pore interaction does not explain the observed exclusion of retained introns from the pore. For example, *FUS* has 10 other introns that appear fully spliced (and presumably loaded with EJCs); yet the presence of 2 retained introns is sufficient to exclude these transcripts from the pore — a finding made possible by the coverage and nucleotide resolution of APEX-seq. The nuclear enrichment of retained introns is also observed in our fractionation-seq data (*r* = 0.78); over 85% of retained-intron genes identified by nuclear APEX-seq were also found by fractionation-seq.

In addition to retained introns, we identify a family of RNAs that show no gene-level subcellular localization differences but exhibit substantial spatial dynamics at the transcript isoform level between the nucleus and cytoplasm (“isoform-switching”, Figure 4E, S5A-E). Thus, the apparent ubiquity of an RNA in two locations is due to the overlay of two isoforms of the RNA: one isoform in location *A* and another isoform in location *B*. Examples of isoform switching localization include mRNAs of the oncogene *AKT2* and circadian rhythm gene *CSNK1D* (Figure 4E). While some of the isoform-switching transcripts have retained introns, we also find such switching in the 5′ UTR, 3′ UTR and coding regions of transcripts (Figure 4F-H). For example, KAT2A (lysine acetyltransferase 2A) shows an isoform with a longer exon in the nucleus. Similarly, NCBP3 (nuclear cap binding subunit 3), an RNA-binding protein contributing to RNA export, shows a transcript with a longer 5′ UTR in the nucleus. Likewise, HNRNPU (heterogenous nuclear ribonucleoprotein U), a DNA- and RNA-binding protein that has many different metabolic functions, expresses a lowly-abundant nuclear isoform with a longer 3′ UTR. Overall we find hundreds of genes with alternative 5′ and 3′ splice sites in our data (Figure S5F-G, FDR < 0.05). These results naturally nominate specific exons associated with each isoform for localization to specific subcellular locations, which in turn could affect downstream functions(Berkovits and Mayr, 2015). Furthermore, our data could be useful for identifying neoepitopes in cancers resulting from intron retention, as recently reported(Smart et al., 2018). More generally, these results highlight the advantage of APEX-seq in providing full length transcript data, not merely gene identity. Isoform switching would likely be missed and multiple isoforms averaged together in imaging methods based on short probes(Chen et al., 2015b),(Shah et al., 2016).

### m6A Modification and RNA Length in Nuclear Pore Localization

Our findings from observing nuclear-retained introns (Figure 4A-D) confirm that a majority of transcripts enriched at the nuclear-facing nuclear-pore are processed, and thus ready for nuclear export. Previous studies have implicated a variety of proteins in the export process(Katahira, 2015). While the processing of nuclear export is complex and highly-regulated, it is thought that the rate-limiting step for mRNA transport is not transition through the central channel, but rather access to and release from the NPC, at least for the small number of transcripts studied in detail(Grunwald and Singer, 2010; Ma et al., 2013). Furthermore, only about one third of mRNAs entering the NPC complete their export, whereas the remaining abort their journey(Wickramasinghe and Laskey, 2015). *N6*-methyladenosine (m6A) modification of pre-messenger RNAs has been reported as a “fast track” signal for nuclear export(Roundtree et al., 2017). Moreover, RNA length has been hypothesized as a feature influencing RNA export. Long RNAs, corresponding to larger RNP complexes, are thought to take more time to remodel to pass through the nuclear pore. Hence, intuition would suggest that long RNAs will have a corresponding longer dwell time at the inner face of the nuclear pore(Grunwald et al., 2011), leading to increased nuclear pore APEX-seq enrichment. Conversely, when mRNAs are assembled into mRNPs, shorter RNAs may be associated with fewer RBPs, including those necessary for recognition and passage through the nuclear pore.

Because APEX-seq represents a snapshot of the abundance of RNAs at a locale during a short time window, enrichment in nuclear pore APEX-seq should be inversely proportional to the “dwell time” of the RNA at the NPC. If a RNA species transits through the NPC very quickly, fewer molecules will be just at NPC during the labeling window in a population of asynchronous cells, leading to low APEX-seq enrichment. Conversely, if a RNA species requires a prolong delay at the NPC before its passage, then more molecules will queue up at the NPC on average in a population of cells, leading to high APEX-seq enrichment. First, intersection of nuclear-pore APEX-seq data with m^6^ A modification sites in HEK293 cells(Meyer et al., 2012) showed a significant depletion of m^6^ A in transcripts enriched by nuclear pore, compared to nuclear lamina or the cytosol (Figure 4I, p = 2.4 × 10^−5^ and 1.5 × 10^−7^ respectively). This result is consistent with the recently discovery of m6A facilitating nuclear export(Roundtree et al., 2017), and suggests that unmodified RNA transcripts have a longer dwell time at the NPC. Second, when we examine RNA length in our nuclear-pore APEX-seq data, we find the transcripts enriched at the pore in fact tend to be shorter than transcripts at other nuclear locations, contrary to the prevailing model. The inverse relationship between RNA length and nuclear pore APEX enrichment is significant both in the mature transcript (*p* = 4.2 × 10^−4^, Figure 4K and S5H) and the introns length (*p* = 3.7 × 10^−9^, Figure 4J and S5K). For protein-coding transcripts, this difference is most significant at the 3′ UTRs (Figure 4K). However, we find no significant difference in the number of introns among enriched transcripts in the various nuclear locations. Although there exist different processes for export of intronless mRNAs(Delaleau and Borden, 2015), such as recruitment of hnRNPC(McCloskey et al., 2012), we did not observe a significant difference in the proportion of intronless-transcripts at the pore relative to other locations (Figure S5I).

Why might shorter transcripts be preferentially enriched at the nuclear pore? If shorter transcripts took more time to be transported through the pore, we might enrich for such transcripts by APEX-seq. Such a mechanism would tend to even out the transport rate of long and short RNAs through the nuclear pore, potentially reducing the challenge of matching the kinetics of mRNA export and gene expression, e.g. for multi-protein complexes with subunits encoded with drastically different mRNA lengths. A recent cell fractionation study on RNA nuclear export rates in *Drosophila* cells reported a negative correlation between 3′ UTR length nuclear export(Chen and van Steensel, 2017). While these results in *Drosophila* cells may at first appear inconsistent with our findings in human cells, the commonality of the 3’UTR being highlighted is worth noting. The authors further speculate that the binding of regulatory proteins to the RNA may slow down export or retain transcripts in the nucleus. Given the many proteins involved in nuclear export further studies are needed to tease out cause and effect. Nonetheless, our data raise the intriguing hypothesis that an inverse relationship exists between RNA length and nuclear-pore dwell time.

### RNA Repeats and Genomic Position Correlate with Nuclear RNA Localization

Deeper analysis of APEX-seq data suggested further principles of RNA localization within the nucleus. First, we identified specific sequence repeats(Hubley et al., 2016) in the exons of transcripts located in distinct nuclear landmarks (Figure 5A). Repeat sequences make up a majority of the human genome(de Koning et al., 2011), with interspersed nuclear elements SINE (short) and LINE (long) containing retrotransposable (transposable via RNA intermediates) elements that can be deleterious when active and randomly moving to new sites in the genome(Ichiyanagi, 2013). We observed enrichment of SINEs and LINEs within the different nuclear locations (Figure 5A, S6A-D), with the highest enrichment of these elements in the nuclear lamina. The cytosolic locations and the nuclear pore showed no enrichment (Figure S6E).

**Figure 5:**
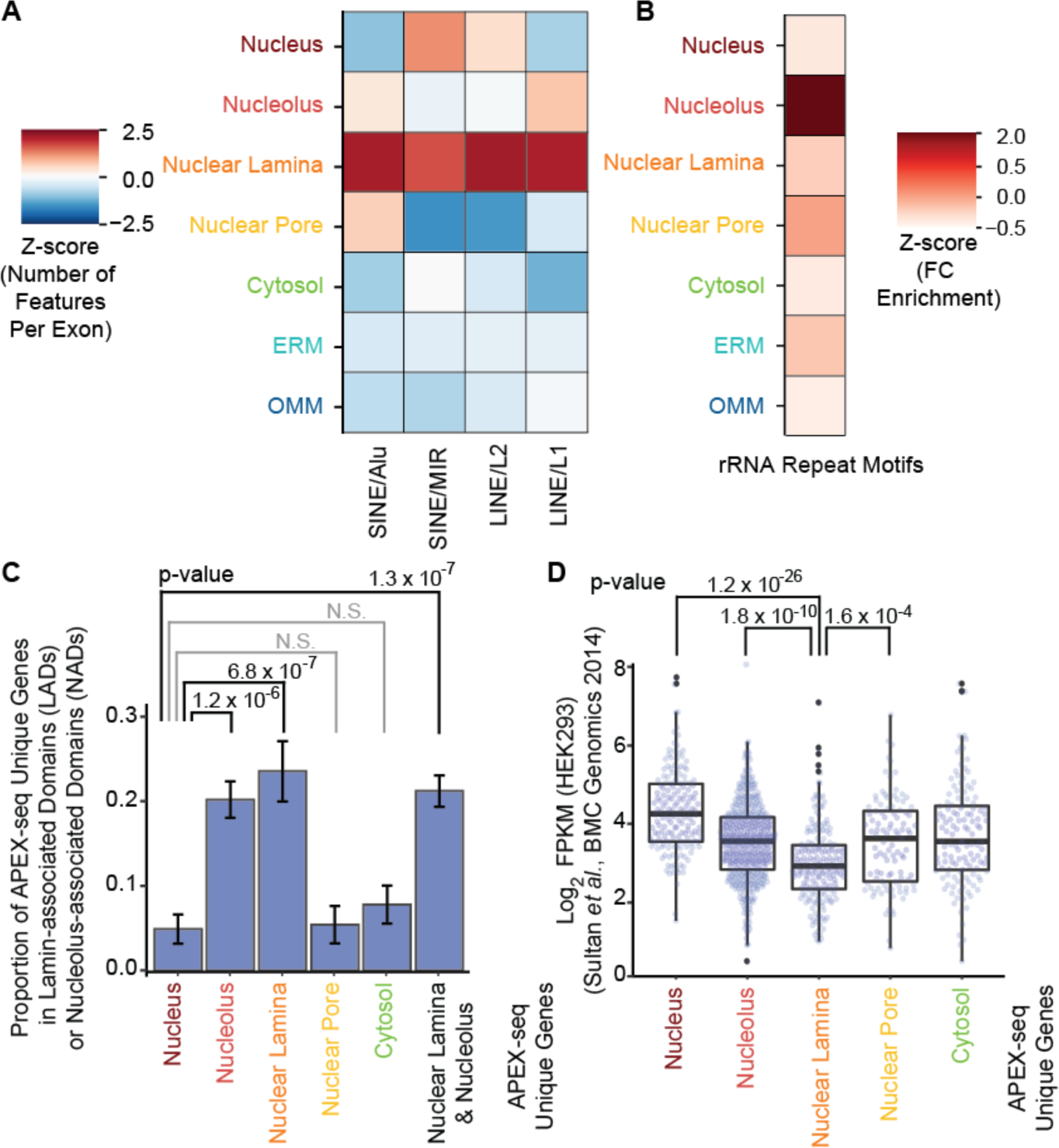
The underlying features of nuclear RNA localization. (A) Examination of repeat elements (including the retrotransposable elements LINE/L1 and LINE/L2) in transcripts uniquely localizing to different locations show an enrichment of these elements in the nuclear-lamina transcriptome. SINE/Alu and SINE/MIR have recently been shown to localize to the nucleus, and here we show these motifs are specifically enriched in the nuclear lamina RNAs. Heatmap scale is the z-score. Figure S6A-D shows the statistics supporting this figure, as well as the heatmap (Figure S6E) obtained when considering all enriched genes, not just uniquely-enriched genes. (B) Heatmap of z-score showing that transcripts localizing to the nucleolus are enriched in rRNA repeat motifs, relative to the nucleus. This finding is consistent with the hypothesis that DNA localization within the nucleus at least partially dictates RNA localization. (C) Examination of the genes found in DNA lamina-associated domains (LADs) and nucleolus associated domains (NADs) confirms that the corresponding transcriptomes are enriched for those genes. Here we restrict to transcripts uniquely enriched in the respective locations. We again show that DNA localization predicts RNA localization within the nucleus. Previous experiments have suggested that NAD and LAD genes have significant overlap(van Steensel and Belmont, 2017). P-values are from Fisher’s exact tests. Figure S6J-M show analysis of NADs and LADs carried out separately for all enriched RNAs (not just uniquely-enriched RNAs), as well as an appropriate control. (D) Within the nuclear locations, the unique nuclear-lamina-enriched transcripts have a lower abundance relative to both the nucleus and the nucleolus. This observation is consistent with the hypothesis that within the nucleus DNA localization partly explains the corresponding RNA localization, as the DNA of genes that tend to be lowly-expressed or silenced are brought to the nuclear lamina. P-value is from a Mann-Whitney U test, FPKM data from Sultan *et al.*(Sultan et al., 2014). Figure S6G-I show nuclear-lamina transcripts have lower abundance when considering all genes, not only genes uniquely enriched in the location.

A recent study has suggested that LINE/L1 elements are epigenetically silenced(Liu et al., 2018), and our finding of these transcripts at the nuclear lamina, where heterochromatin resides, is consistent with this notion(Padeken et al., 2015). Importantly, an RNA sequence derived from SINE/Alu elements was recently shown to be sufficient to drive nuclear retention of some long non-coding RNAs(Chen, 2018; Lubelsky and Ulitsky, 2018), thus suggesting a pathway for nuclear accumulation of both lncRNAs and mRNAs that contain such sequences. Furthermore, removal of repeat sequences from some nuclear lincRNAs has been shown to cause them to relocalize to the cytosol(Carlevaro-Fita et al., 2017). Our APEX-seq data are consistent with these previous studies and further suggest that SINE/Alu-associated nuclear localization is due to preferential localization of these transposable-derived RNAs to the nuclear lamina.

Second, location of the DNA locus from which an RNA originates strongly dictates nuclear RNA location. The genome is hierarchically organized with specific classes of DNA elements residing in distinct nuclear domains and bodies(Dekker et al., 2017). For example, the nucleolus is enriched for DNA coding for ribosomal RNAs (rRNAs)(van Koningsbruggen et al., 2010), and rRNA repeat motifs(Wheeler et al., 2013) (typical size ~10^2^-10^3^ bp) are highly enriched in the nucleolus by APEX-seq, but far less so in the nuclear lamina or cytosol (Figure 5B). Beyond rRNAs, mRNA of genes residing in DNA nucleolus-associated domains (NADs)(Dillinger et al., 2017; van Koningsbruggen et al., 2010) are similarly enriched in the nucleolus by APEX-seq (*p* = 4.9 × 10^−4^; odds ratio = 4.4; 95% confidence interval (CI) = 1.7 – 14) (Figure 5C, S6J). Conversely for DNA loci in nuclear lamina-associated domains (LADs)(Guelen et al., 2008), their corresponding RNA were enriched in the lamina APEX-seq (*p* = 2.2 × 10^−8^; odds ratio = 11; 95% CI = 3.8 – 43) (Figure S6J-M). Transcripts enriched at the nuclear lamina also had lower expression level than those in the nucleus in general (*p* = 1.2 × 10^−26^), or transcripts at the nucleolus (*p* = 1.8 × 10^−10^) or nuclear pore (*p* = 1.6 × 10^−4^), consistent with the idea of heterochromatin deposition and gene silencing at LADs (Figure 5D, S6G-I). Neither the nucleolus nor nuclear lamina were enriched in mitochondrial genes, which we do not expect to localize within a specific nuclear sub-compartment (Figure S6M). Taken together, a strong relationship between the mature (polyadenylated) RNA organization and the underlying genome organization emerges, with the nucleolus and nuclear lamina defining a radial axis for nuclear RNA positions.

### Distinct Mechanisms for mRNA Localization to the Outer Mitochondrial Membrane

The human mitochondrion contains >1100 distinct protein species(Calvo et al., 2015), only 13 of which are encoded by the mitochondrial genome (mtDNA) and translated within the mitochondrion. The remainder are encoded by the nuclear genome (nDNA) and must be delivered to the mitochondrion after translation in the cytosol(Mercer et al., 2011). The identification of ribosomes at the outer mitochondrial membrane (OMM) by electron microscopy decades ago(Kellems et al., 1974, 1975) led to the hypothesis that some mRNAs encoding mitochondrial proteins may be locally translated at the OMM and post-translationally or co-translationally delivered into the mitochondrion(Gold et al., 2017). Recently, proximity-specific ribosome profiling(Ingolia et al., 2009) revealed that, in yeast, a subset of mRNAs encoding mitochondrial proteins are indeed translated locally at the OMM(Williams et al., 2014). However, a similar study could not be performed in mammalian cells, because the biotin starvation required for achieving spatially specific biotinylation in ribosome profiling is toxic to mammalian cells. Purification of mammalian mitochondria followed by RNA-seq has also failed to identify OMM-associated mRNAs, likely because the centrifugation steps required for mitochondrial fractionation disrupt the OMM and RNA binding interactions there(Lesnik et al., 2015; Mercer et al., 2011). Hence, at present, we know very little about the landscape of RNAs at the mammalian mitochondrial membrane, despite the importance of this information for understanding the mechanisms of mitochondrial biogenesis.

Our APEX-seq data revealed strong enrichment (Z-score >2) of mRNAs encoding mitochondrial proteins at the outer mitochondrial membrane compared to the 8 other compartments that we tested (Figure 3J). We also plotted the OMM APEX-seq data by enrichment score and observed a significant right-shift in nuclear-encoded mitochondrial genes (red) over non-mitochondrial/non-secretory genes (Figure 6A; p <10^−50^, Mann-Whitney U test). By contrast, this enrichment of mitochondrial genes is not observed in the ERM APEX-seq dataset (Figure 6B). Examination of the OMM-enriched mRNA population did not reveal any pattern in terms of protein functional class or sub-mitochondrial localization of the encoded proteins. We reasoned that under basal conditions, multiple mRNA subpopulations that are targeted by different mechanisms to the OMM may overlap and be difficult to distinguish from one another. In an effort to tease apart these possible subpopulations, we repeated OMM APEX-seq labeling under different perturbation conditions.

**Figure 6:**
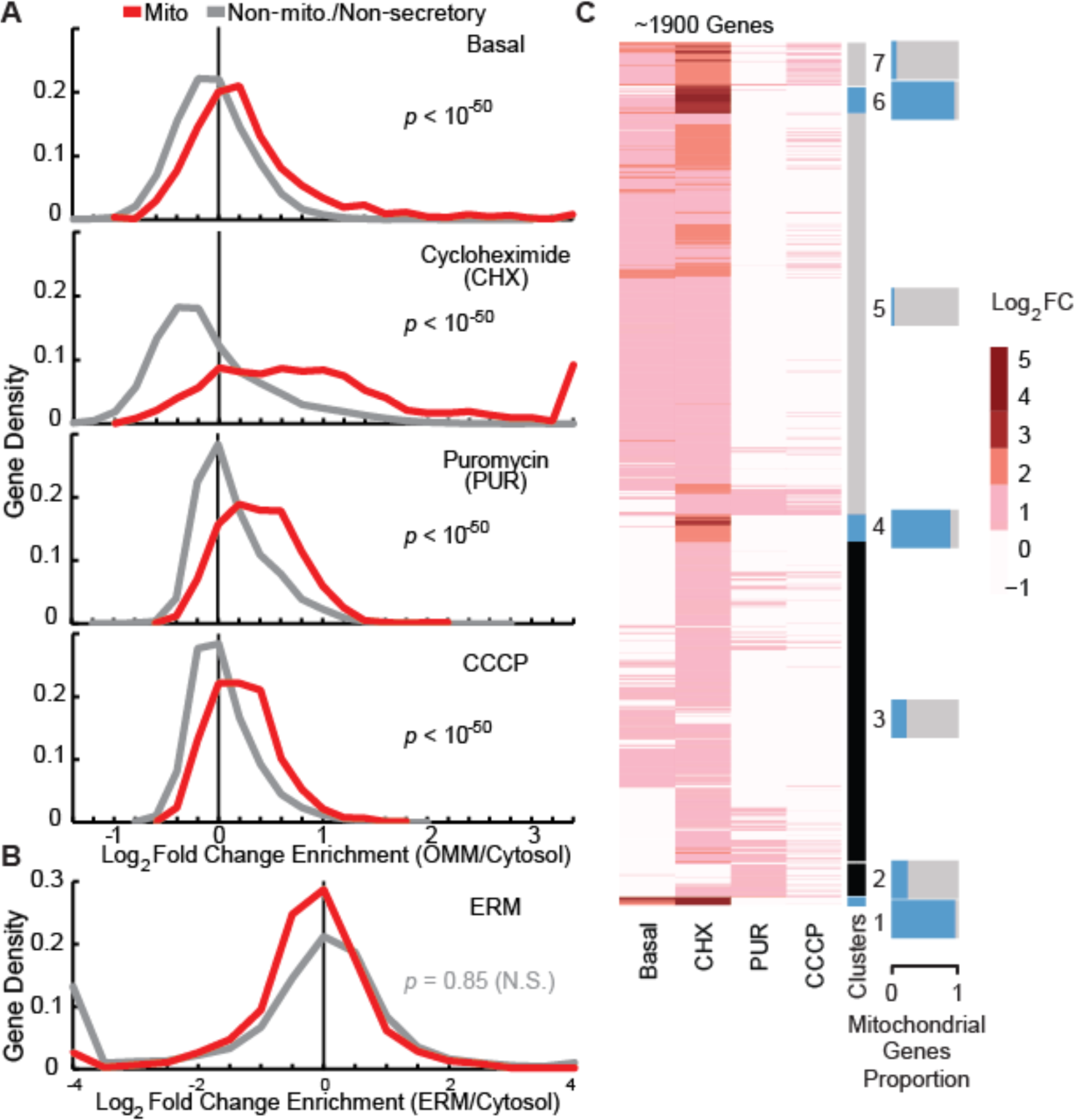
Enrichment of mRNAs encoding mitochondrial proteins at OMM. (A) Gene density distribution of OMM APEX-seq RNA enrichments, in HEK-293T cells under basal, cycloheximide (CHX), puromycin (PUR) or CCCP-treated conditions. P-values comparing the distributions of mitochondrial genes and non-mitochondrial/non-secretory genes are from Mann-Whitney U tests. (B) Gene density distribution of ERM APEX-seq RNA enrichment. Genes are categorized as in (A). P-value comparing the distributions of mitochondrial genes and non-mitochondrial/non-secretory genes is from a Mann-Whitney U test. (C) Heatmap of the fold changes for transcripts enriched by OMM APEX-seq. Upon clustering based on the basal, cycloheximide (CHX) and puromycin (PUR) experiment, we obtain clusters of transcripts that are either strongly enriched or depleted in the corresponding mitochondrial proteins. Figure S7B shows the corresponding correlation plots. Figure S7C shows GO-terms associated with the clusters.

Taking advantage of the rapidity of APEX-seq tagging, we treated HEK cells expressing OMM-APEX2 with cycloheximide (CHX), puromycin (PUR), or carbonyl cyanide m-chlorophenyl hydrazone (CCCP), prior to biotin labeling (Figure S7A; correlation *r* of replicates between 0.97 and 1). CHX (also used for proximity-specific ribosome profiling(Williams et al., 2014)) and PUR are both protein translation inhibitors but they work by different mechanisms; CHX stalls translation but preserves the mRNA-ribosome-nascent protein chain complex, while PUR dissociates mRNAs from ribosomes. CCCP is a drug that abolishes the mitochondrial membrane potential and thereby stops membrane potential-dependent processes including TOM (translocase of outer membrane)/TIM-mediated import of mitochondrial proteins(Chacinska et al., 2009).

After treatment of cells with CHX, we observe a dramatic increase in both the number of mitochondrial genes and their extent of enrichment at the OMM (Figures 6A and 7A). As noted in the yeast OMM ribosome profiling study(Williams et al., 2014), CHX likely increases the interaction between nascent peptide chain-ribosome-mRNA complexes and the TOM complex on the OMM that recognizes mitochondrial targeting peptides(Chacinska et al., 2009). Further analysis showed that the top-most enriched mitochondrial genes under the CHX condition have higher TargetP scores on average (Figure 7B-C), an indicator of the protein product’s mitochondrial targeting potential. Proteins with high TargetP include soluble mitochondrial matrix-resident proteins involved in TCA cycle, amino acid metabolism, and mtDNA functions (Figure 7E-F). Hence, OMM APEX-seq following CHX treatment appears to highlight a population of high TargetP mitochondrial mRNAs that may localize to the OMM in a ribosome-dependent fashion, possibly via interactions between nascent chain and TOM (Figure 7I). Figure 7D shows the genome track of *HSPA9* (mitochondria heat shock protein A9, with targetP score of 5), with increased localization of the RNA to the OMM upon CHX treatment; *MUT* (methylmalonyl-coA mutase) (Figure S7G) is another RNA showing similar localization.

**Figure 7:**
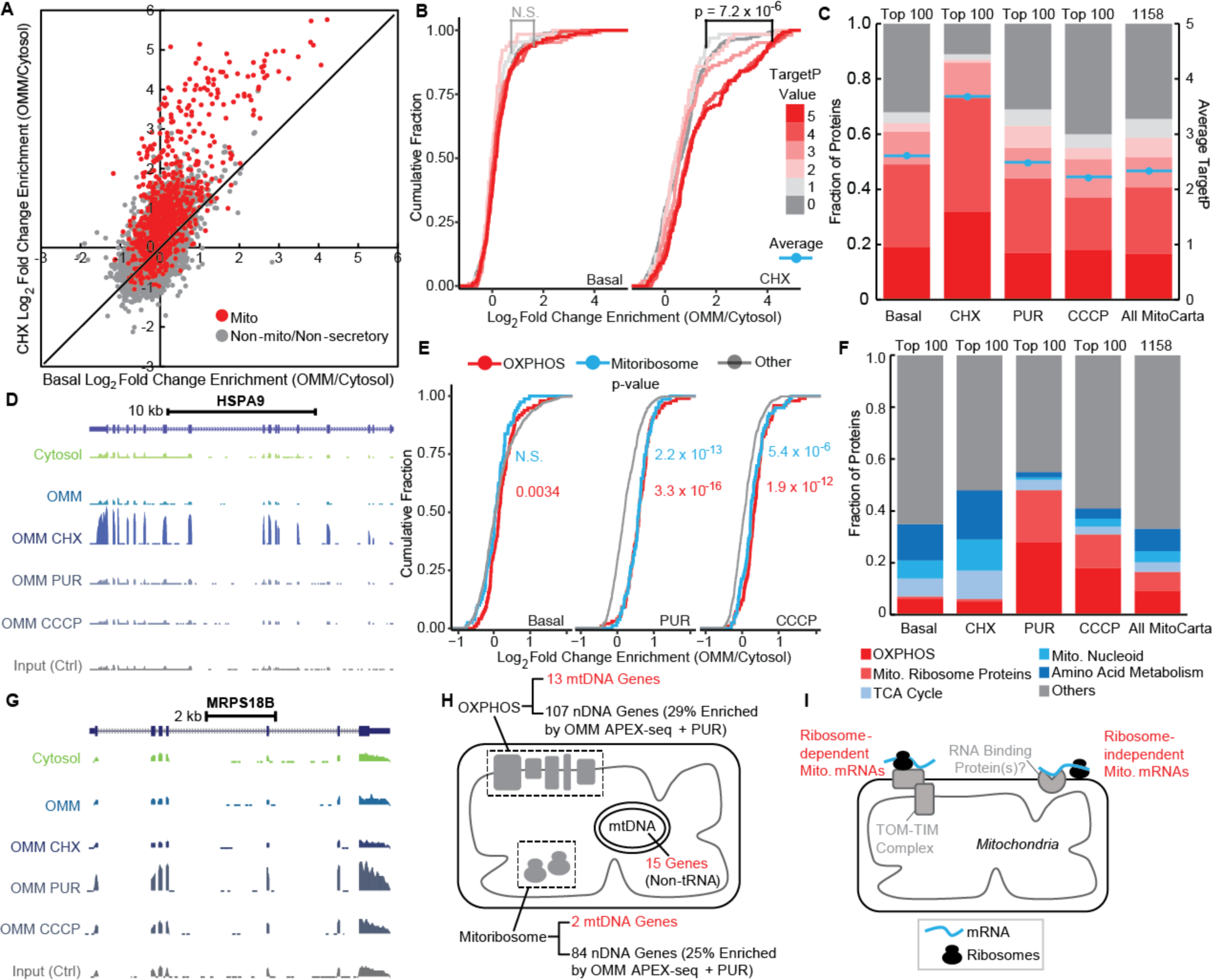
Distinct subpopulations of mRNAs at the OMM revealed by APEX-seq. (A) Scatter plot of OMM APEX-seq log_2_ fold-change in HEK-293T cells comparing the basal (x axis) and CHX (y axis) conditions. Genes are categorized as in Figure 6A. (B) Cumulative fraction of genes in different conditions by TargetP values. CHX treatment shows increased OMM targeting of genes with high Target P values. Genes are categorized by their TargetP values (adapted from MitoCarta; see Methods section) on a scale from 5 (strongest N-terminal mitochondrial targeting peptide) to 0 (no N-terminal mitochondrial targeting peptide). (C) Comparing the proportion of transcripts with different TargetP values (y axis, left) and average TargetP value (y axis, right) among top 100 mitochondrial genes enriched by OMM APEX-seq in HEK-293T cells under CHX, basal, PUR, CCCP conditions, and all 1158 MitoCarta genes (right-most bar). (D) UCSC Browser tracks of a mitochondrial gene (*HSPA9*, targetP=5) show increased enrichment by OMM-APEX upon CHX treatment. Figure S7G shows the track from another RNA *MUT* (methylmalonyl-coA mutase) that shows increased enrichment upon CHX treatment. (E) Cumulative fraction of OXPHOS and mitoribosome related genes in different conditions. Both PUR and CCCP treatment show increased OMM targeting of OXPHOS and mitoribosome related genes. Genes are functionally classified according to Gene Ontology. (F) Comparing the proportion of transcripts in different functional classes among top 100 mitochondrial genes enriched by OMM APEX-seq in HEK-293T cells under, basal, CHX, PUR, CCCP conditions, and all MitoCarta genes. Genes are functionally classified according to Gene Ontology. (G) UCSC Browser tracks of a mitochondrial ribosomal gene (*MRPS18B*) that show increased enrichment by OMM-APEX upon PUR/CCCP treatment. (H) Scheme illustrating the coordinated assembly of respiratory chain complexes and mitoribosomes between the nuclear genome and the mitochondrial genome. (I) Model summarizing two distinct subpopulations (red-labeled groups) of mitochondrial RNAs proximal to mitochondria.

Treatment of cells with PUR produced a pattern of enrichment distinct from CHX treatment. PUR dissociates mRNAs from ribosomes and nascent chains, and the vast majority of CHX-enriched mRNAs are no longer observed at the OMM, consistent with the hypothesis that the localization of these transcripts depends on an intact ribosome complex (Figure 6A). Nonetheless, a subpopulation of mRNAs remained clearly associated with the OMM (Figure 6A). The top PUR-enriched genes are not higher in TargetP, in contrast to CHX-enriched genes (Figure 7C). Functional class analysis reveals that PUR-enriched genes have a higher likelihood of encoding mitochondrial ribosome and oxidative phosphorylation (OXPHOS) components (Figure 7E-F); Figure 7G shows genome tracks(Kent et al., 2002) of a representative mitochondrial ribosomal protein gene, *MRPS18B* (28S ribosomal protein S18b), whereas Figure S7H shows the track of a representative OXPHOS gene, *NDUFB9* (NADH:ubiquinone oxidoreductase subunit B9). These trends may not have been apparent under basal conditions due to overlap with and obscuring of signal by other mRNA populations. The PUR data suggest that a subpopulation of mRNAs associates with the OMM in a ribosome and nascent chain-independent fashion, perhaps by binding directly to an RNA binding protein localized to the OMM (Figure 7I).

Why might this ribosome-independent OMM-proximal mRNA population be specifically enriched for mitochondrial ribosome and OXPHOS genes? One possibility is for the purpose of coordinating their translation with intra-mitochondrial translation and transcription. All 15 of the non-tRNA genes in mtDNA code for mitochondrial ribosome and OXPHOS components (Figure 7H). Previous work using mitochondrial and cytosolic ribosome profiling showed that intra-mitochondrial translation and cytosolic translation of mitochondrial proteins are intimately coordinated in yeast(Couvillion et al., 2016) and likely in human cells(Richter-Dennerlein et al., 2016). Perhaps one mechanism for achieving this coordination is to localize OXPHOS and mitochondrial ribosome mRNAs to the OMM via a specific RNA binding protein that senses intra-mitochondrial translation status.

Upon treatment with CCCP, the mitochondrial genes enriched at the OMM are similar to PUR-enriched genes (Figure 7C, 7E and 7F), indicating that their OMM localization does not depend on the mitochondrial membrane potential. We speculate that because CCCP inhibits mitochondrial protein import, this decreases interactions of ribosome-mRNA-nascent chain complexes with the OMM. Under these conditions, association of ribosome-independent mRNAs (the same ones enriched by PUR) with the OMM becomes clearer.

The availability of the basal OMM APEX-Seq dataset along with three “drug perturbation” OMM APEX-Seq datasets enabled us to perform higher order clustering analysis. Figure 6C shows transcripts that were enriched at the OMM in at least one condition (log_2_ fold-change > 0.75, *q* < 0.05). We find that RNAs cluster into groups based on their enrichment factors in CHX versus PUR; the effects of CCCP, which are correlated with the effects of PUR (*r*_pearson_ = 0.53) are shown for comparison. We found some clusters of enriched transcripts to be strongly predictive of genes coding for mitochondrial proteins (Figure S7D). In particular, in clusters 1, 4, and 6 that included transcripts strongly enriched upon CHX treatment and depleted upon PUR treatment, > 90% of RNAs (N = 128/140) code for mitochondrial proteins. While 7 of the remaining 12 transcripts were pseudogenes, of the 5 mRNAs that were in those clusters but not in MitoCarta 2.0, at least 3 are likely to be mitochondrial (Figure S7E-F) based on other studies(Mou et al., 2009; Pandey et al., 2017; Thul et al., 2017). Similarly, clusters 5+7 that were depleted upon PUR treatment but not enriched by CHX were strongly depleted for mitochondrial genes (<5%, N = 45/990). Thus, OMM APEX-seq data could be used to predict whether certain genes will code for mitochondrial proteins.

## DISCUSSION

APEX-seq is a powerful proximity sequencing technology for RNA. APEX-seq yields complete RNA sequence information to single nucleotide resolution, thereby filling a critical gap in the landscape of RNA technologies. Our map of transcriptome localization provides one of the most comprehensive and precise delineations of RNA spatial organization in the living cell. We highlight patterns and principles of RNA localization related to its birth, primary sequence, isoform processing, and ultimate fates of the encoded proteins.

With quantitative enrichment scores and detailed transcript profiles for over 25,000 distinct human RNA species across 9 subcellular compartments, our study reveals new patterns of RNA localization that give rise to a variety of biological hypotheses. We speculate on the role of the nuclear pore in gating RNA export and enriching shorter transcripts, on the extensive overlap between mitochondrial membrane and ER membrane-associated transcriptomes, on the pervasive diversity in localization of different transcript isoforms, on the ability of RNA repeats and genomic position to shape nuclear RNA organization, and on distinct mechanisms for mRNA targeting to the mitochondrial outer membrane.

APEX-seq adds to arsenal of RNA localization methods, and offers unique advantages and disadvantages compared to existing techniques. The first strength of APEX-seq is that labeling is performed in living cells, while all membranes and macromolecular complexes are still intact. The features enables APEX-seq to probe “unpurifiable” structures such as the nuclear lamina (that cannot be accessed by fractionation-seq, for example), and also to achieve higher specificity in compartments that can be purified, such as the nucleus. Live cell labeling also circumvents artifacts associated with cell fixation, as required for imaging-based methods such as MERFISH and in-situ sequencing. The second strength of APEX-seq is that it provides full sequence information for diverse classes of RNA transcripts, allowing transcript isoforms with distinct localization to be readily distinguished (Figure 4F-H) – a task that would be challenging or impossible to do on a transcriptome-wide scale using short probe-based imaging methods. In contrast to ribosome profiling, which captures actively translating mRNA on polysomes, APEX-seq additionally detects lncRNAs, antisense RNAs (Figure 3C-D) and untranslated mRNAs not bound by ribosomes, as seen in the OMM APEX-seq puromycin perturbation experiment (Figure 6). Thirdly, the high spatiotemporal resolution (1-minute direct RNA labeling and nanometer spatial specificity) sets APEX-seq apart from APEX-RIP, which suffers from low temporal resolution due to long time required for formaldehyde crosslinking (>17 minutes) and poor spatial specificity in non-membrane enclosed regions of the cell (Figure 1D).

A disadvantage of APEX-seq is that it requires an APEX fusion construct to be recombinantly expressed in the sample of interest, which limits applicability to human tissue, for example. Also, APEX-seq is performed on populations of cells and does not provide single-cell information like imaging-based methods. Finally, because labeling is performed in live cells, APEX-seq coverage will be fundamentally limited by the steric accessibility of RNAs in their native context; RNAs that are buried within macromolecular complexes may not be tagged and enriched by APEX-seq whereas they could be detected by lysis and purification-based methods. These limitations suggest directions for future improvement.

We expect that APEX-seq will be broadly applicable to many organisms and cell types, just as APEX proteomics has been extended to flies(Chen et al., 2015a), worms(Reinke et al., 2017), yeast(Hwang and Espenshade, 2016), and neurons(Loh et al., 2016). APEX-seq could be fruitfully applied to polarized cells, cells with long extensions (e.g. neurons), or dynamic developmental systems. Future use of APEX-seq in conjunction with RNA structure mapping methods(Chin and Lecuyer, 2017; Spitale et al., 2015), RBP occupancy atlases(Van Nostrand et al., 2016), and massively parallel reporter gene assays(Lubelsky and Ulitsky, 2018; Shukla et al., 2018) could shed light on the molecular basis of the exquisite and extensive spatial organization of RNA within the cell.

## ACKNOWLEDGMENTS

We thank members of the Ting and Chang labs for helpful discussions. We thank J. He, P.J. Batista and M.R. Corces for analysis suggestions; J. Coller and X. Ji for sequencing; and L. Mateo for assistance with microscopy. We thank A.C. Carter and B.S. Zhao for critical reading of the manuscript.

## Funding

This work was supported by NIH (R01-CA186568-1 to A.Y.T.; R01-HG004361 and P50-HG007735 to H.Y.C.; S10OD018220 to Stanford Functional Genomics Facility), Burroughs Wellcome (CASI grant to A.N.B.) and the Chan Zuckerberg Biohub (to A.Y.T.). F.M.F. was supported by the NIH T32 Genomics Training Fellowship and the Arnold. O. Beckman Postdoctoral Fellowship. S.H. was supported by the Stanford Bio-X Bowes Fellowship. H.Y.C. is an Investigator of the Howard Hughes Medical Institute.

## Author Contributions

P.K. and A.Y.T. conceived the project. F.M.F., S.H., P.K., H.Y.C. and A.Y.T. designed experiments. F.M.F., S.H. and P.K. performed all experiments, unless otherwise noted. F.M.F. designed and carried out sequencing experiments. S.H., A.N.B. and A.Y.T. designed and carried out sequential FISH experiments. F.M.F, S.H, P.K, K.R.P, J.X. and A.Y.T. analyzed data. F.M.F., S.H., H.Y.C. and A.Y.T. wrote the paper with input from all authors. H.Y.C. and A.Y.T. jointly supervised work.

## Competing interests

A.Y.T., P.K., H.Y.C. and F.M.F. have filed a patent application covering aspects of this work (U.S. Patent Application Number US 2017/0226561). H.Y.C. is a co-founder and advisor of Accent Therapeutics. H.Y.C. is an advisor of 10X Genomics and Spring Discovery.

**Figure S1:**
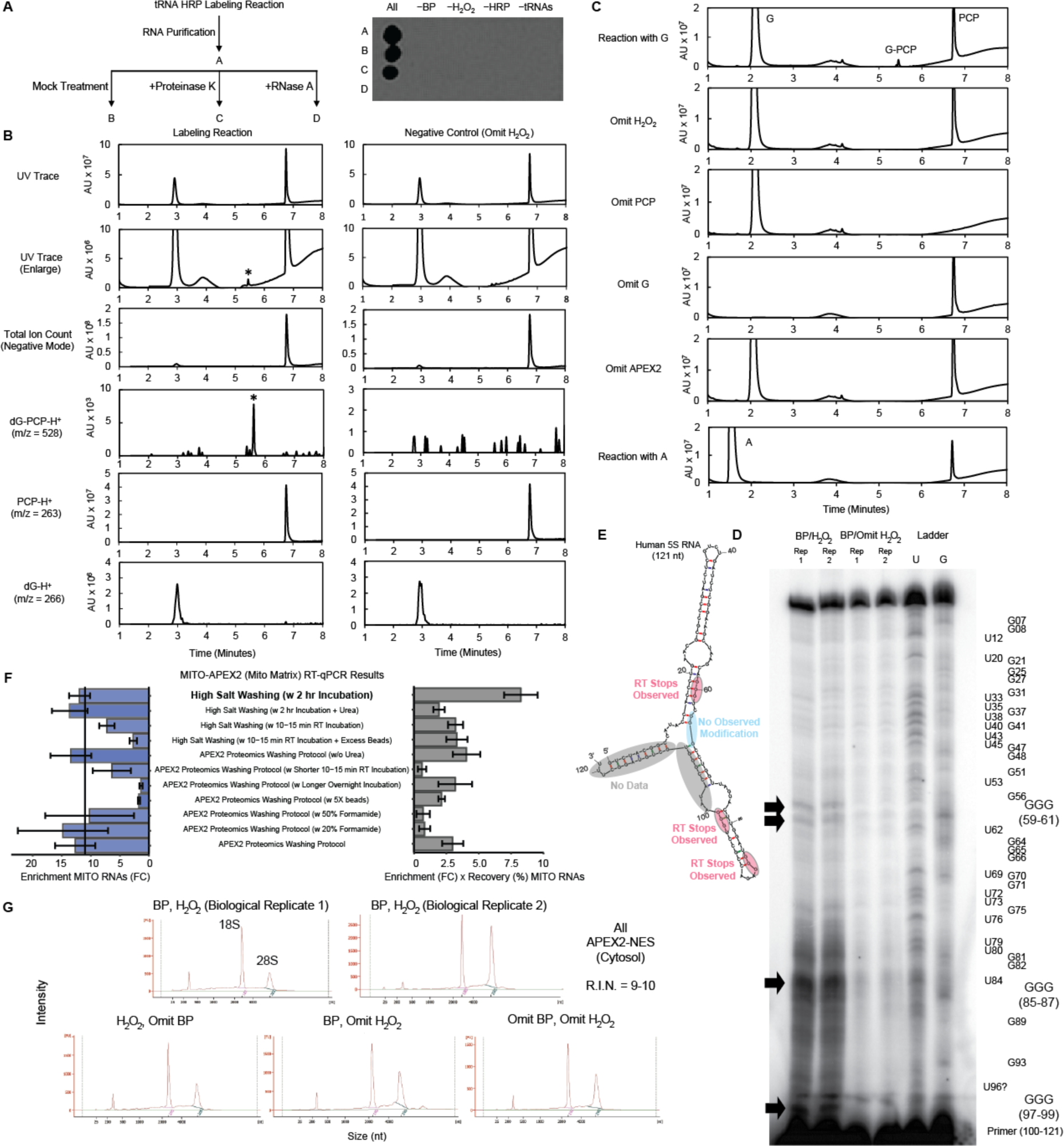
(A) Streptavidin-biotin dot blot analysis of tRNA labeling by horseradish peroxidase (HRP) *in vitro*. Left: Yeast tRNA extract was incubated for 1 minute with HRP, biotin-phenol and H_2_O_2_, after which the tRNA was purified and the resulting product was treated with either proteinase K or RNase A. Right: The products were spotted, and biotinylated species detected via staining with streptavidin. (B) LC-MS detection of deoxyguanosine (dG)-pentachlorophenol (PCP) adduct resulting from HRP labeling reaction *in vitro*. dG was incubated for 1 minute with PCP, HRP, and H_2_O_2_. The chemical composition of the resulting mixture was analyzed by LC-MS in negative ion detection mode. The left column shows the experiment and the right column shows the negative control in which H_2_O_2_ was omitted. Row 1 is the UV trace chromatogram. Row 2 is an enlarged UV trace chromatogram (asterisk denotes the dG-PCP product). Row 3 is the total ion count chromatogram. Rows 4-6 shows mass chromatograms corresponding to the mass of dG-PCP (product), PCP (starting material), and dG (starting material), respectively. (C) APEX2 catalyzes formation of G-PCP adduct. UV trace chromatograms of guanosine (G) or adenosine (A) reacting with PCP and APEX2 *in vitro*. G (top row) or A (bottom row) was incubated for 1 minute with PCP, APEX2, and H_2_O_2_. The resulting mixture was analyzed by HPLC with UV detection of chemical species. Absorption peaks corresponding to G, PCP and G-PCP adduct are labeled in row 1. Rows 2-5 show negative controls with H_2_O_2_, PCP, G or APEX2 omitted, respectively. Row 6 shows the same experiment with A in place of G. (D) *In vitro* transcribed 5S ribosomal RNA was treated with HRP in duplicate in the presence of biotin-phenol followed by 1-minute treatment with H_2_O_2_ and enriched by streptavidin-biotin pulldown. The enriched RNA was reverse transcribed into cDNA, and the resulting products were run on a denaturing PAGE gel. Modification of 5S RNA at GGG sequences results in excess truncated DNA products (black arrows) relative to controls (no H_2_O_2_ added) carried out in duplicate. (E) For 5S RNA, 3 of the 4 GGG sequences interrogated yielded gel bands, presumably due to the RT-enzyme halting at the corresponding biotinylated nucleotides. (F) Optimization of the washing step following binding of biotinylated RNA to the streptavidin beads. We used buffers containing high salt (1M NaCl) as well as buffers previously used for APEX Proteomics washes with or without urea or formamide. We varied both the amount and type of streptavidin beads used, and the duration and temperature at which incubations of labeled RNA with the beads were carried out. A simple high salt wash provided high enrichment of mitochondrial RNAs (blue bar graph), as determined by enrichment by RT-qPCR of two mitochondrial RNAs - *MTND1* and *MTCO1* - relative to 4 non-mitochondrial RNAs - *XIST*, *FAU*, *GAPDH* and *SSR2*). The high-salt wash also maintained higher and more reproducible recovery relative to other conditions. Sources of error arise from both biological replicates and technical replicates. (G) Bioanalyzer traces of RNA extracted from APEX2-NES HEK-293T cells confirming that the RNA is not degraded (RNA integrity number (R.I.N.) = 9-10) upon treatment with biotin phenol and/or hydrogen peroxide, based on the ribosomal RNAs 18S and 28S remaining intact.

**Figure S2:**
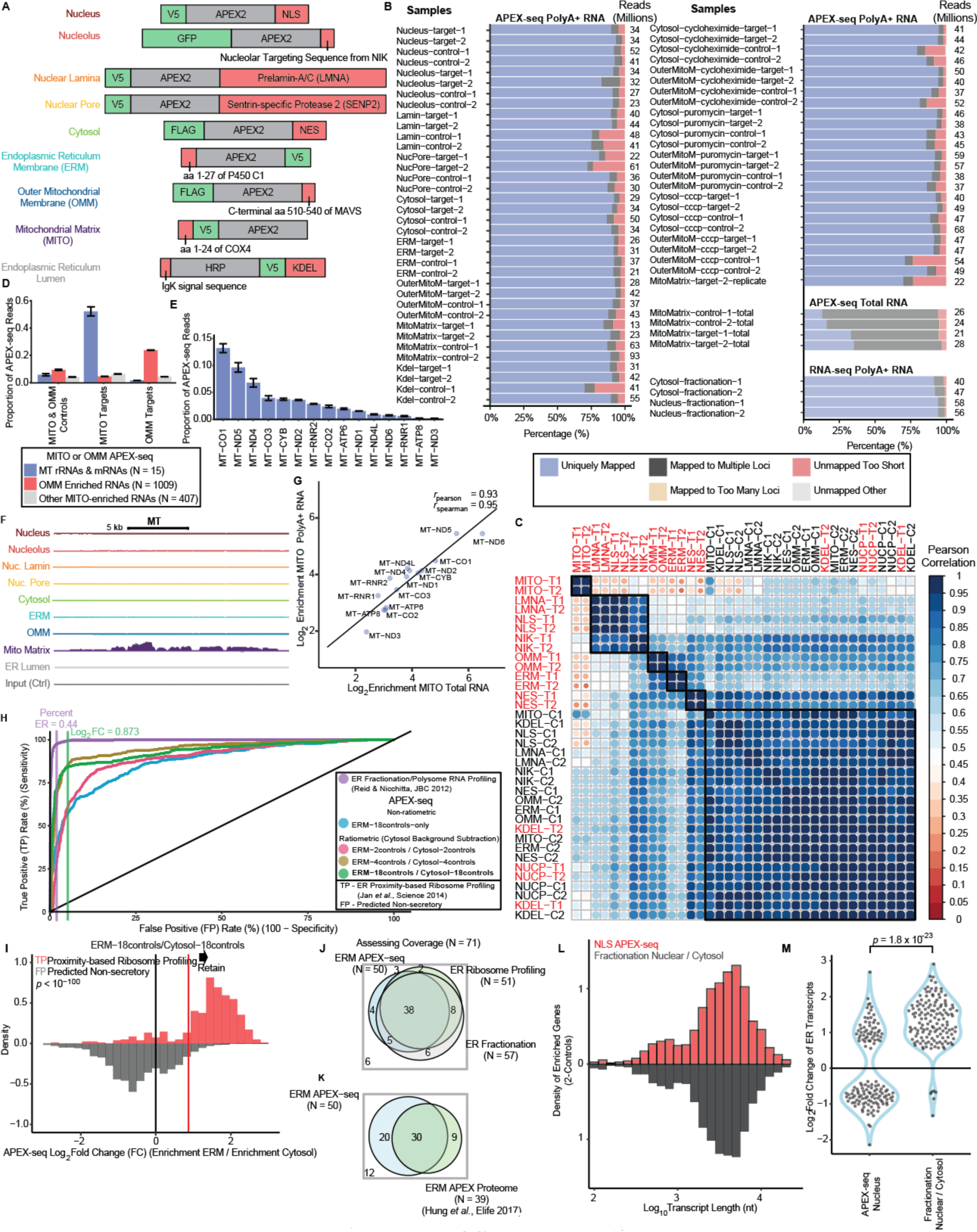
(A) APEX2 fusion constructs employed in this study. FLAG-APEX2-NES uses a nuclear export signal (NES) to localize APEX2 throughout the cytoplasm. Mito-V5-APEX2 employs a 24-amino-acid mitochondrial targeting sequence (MTS) from COX4 to localize APEX2 throughout the mitochondrial matrix. FLAG-OMM-APEX2 employs the C-terminal 31 amino acids of mitochondrial antiviral-signaling protein (MAVS) to target APEX2 to the outer mitochondrial membrane (OMM). ERM-APEX2-V5 employs the transmembrane segment of the endoplasmic reticulum (ER)-resident protein P450 oxidase 2C1 to target APEX2 to the ER membrane (ERM). HRP-V5-KDEL employs a KDEL sequence to target the horseradish peroxidase (HRP) to the ER lumen. V5-APEX2-NLS employs a nuclear localization sequence (NLS) to target APEX2 throughout the entire nucleus. GFP-APEX2-NIK3x employs three tandem nucleolar targeting sequences from NF-κB-inducing kinase (NIK) to localize APEX2 to the nucleolus. V5-APEX2-LMNA is targeted to the nuclear lamina by fusing APEX2 to the N terminus of prelamin-A/C (LMNA). V5-APEX2-SENP2 is targeted to the nuclear pore complex by fusing APEX2 to the N terminus of Sentrin-specific protease 2 (SENP2). V5 and FLAG are epitope tags. (B) Mapping statistics of all APEX-seq and Fractionation RNA-seq libraries generated for this study. Figures show the percentage of mapped reads, as well as the total number of reads. Most polyA**+** libraries showed high proportion (> 80%) of uniquely-mapped reads. APEX-seq libraries showed high mappability, only slightly lower that the corresponding fractionation RNA-seq libraries mapped using the same settings (using the STAR software). (C) Correlation plot of biological replicates, showing that the unlabeled controls for the different constructs are quite similar to each other, and to the nuclear pore and ER lumen target constructs. The MITO APEX-seq libraries are most different from the other libraries. (D) The 15 MT rRNAs and mRNAs (blue) account for more than half of the sequencing reads in the MITO APEX-seq libraries, but ~6% in the control libraries. The corresponding values for OMM-enriched RNAs (red) are also shown, as well as other RNAs (grey) identified as enriched based on the analysis pipeline used. (E) The proportion of all reads in the MITO APEX-seq libraries mapping to the 15 mRNAs and rRNAs. Over 10% of all reads map to *MTCO1*. (F) UCSC genome track of mitochondrial (MT) genome showing robust enrichment of mitochondrial RNAs in the mitochondrial-matrix (MITO) APEX-seq library, but not from the libraries generated from constructs targeting other subcellular locations. (G) Scatter plot of enrichment for the 15 MT rRNAs and mRNAs between polyA+ RNA and total RNA, showing good agreement between the two. (H) ROC curve showing the performance of APEX-seq for different analysis protocols. These include no ratiometric normalization (blue), as well as 2-controls (red), 4-controls (yellow) and 18-controls conditions (green). For challenging open locations, combining controls from other APEX-seq constructs improves performance. For the entire paper, unless otherwise mentioned, 18-controls data is shown. For comparison the ER polysome profiling RNA-seq is shown (purple). Here the true positive is the Jan *et al.* list(Jan et al., 2014), and false positive is predicted non-secretory proteins(Kaewsapsak et al., 2017). (I) ERM APEX-seq shows clear separation of true positives (determined by proximity-based ribosome profiling), and negatives (predicted to be non-secretory based on Phobius, SignalP or TMHMM). (J) Using a list of 71 true-positive ERM transcripts, the coverage of APEX-seq was compared to ribosome profiling and ER fractionation RNA-seq. All three methods yield similar coverage. (K) Comparing ERM APEX-seq to ERM proteomics, APEX-seq shows higher coverage for these 71 genes. (L) Histogram showing the length distribution of transcripts recovered from both methods. Both methods yield comparable distributions for the length of transcripts recovered. (M) Violin plot showing the striking difference in fold changes of ER transcripts between fractionation and APEX-seq. P-value is from a Mann-Whitney U test.

**Figure S3:**
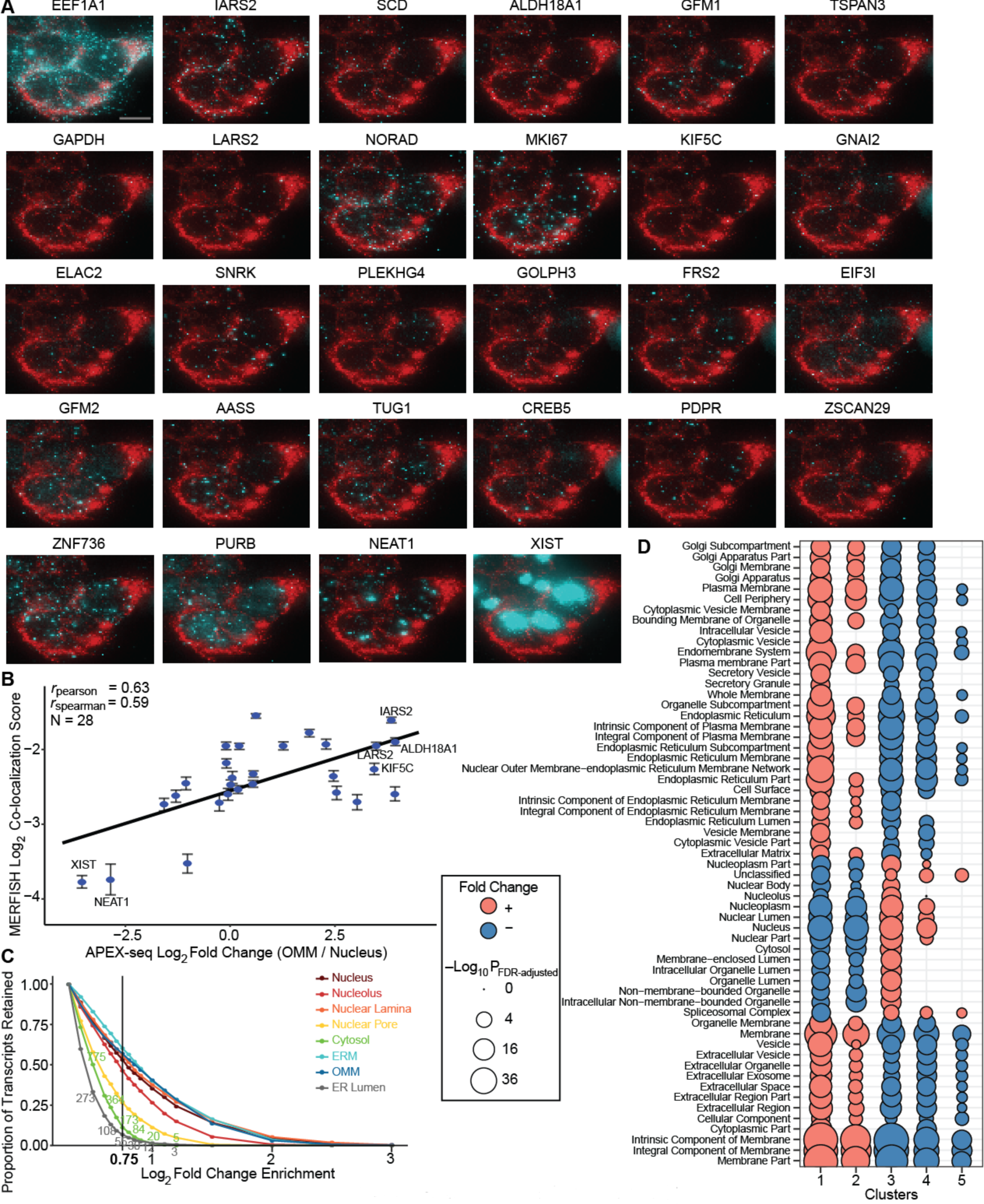
(A) Imaging the spatial localization of 28 selected genes in HEK-293T cells using sequential FISH. Each individual gene (cyan, pseudocolor) is shown together with *MTND3* (red, pseudocolor) as a marker. Panels are arranged by decreasing co-localization with *MTND3*. Scale bar, 10 μm. (B) Correlation of FISH co-localization score versus APEX Seq OMM/nucleus fold change. FISH co-localization score was calculated as the percentage of total signal overlapping with a mitochondrial mRNA, *MTND3*. (C) The proportion of transcripts retained as enriched, as the fold-change cutoff based on APEX-seq comparison of labeled targets versus unlabeled controls is varied. Unless otherwise mentioned, a log_2_ fold-change of 0.75 was used in Figure 2 and 3. (D) Cellular component GO-terms associated with the clusters determined from heatmap in Figure 3B, confirming that the nuclear locations are enriched for nuclear-associated GO terms, and the ER and OMM for membrane GO terms. Size of bubble denotes more significant enrichment/depletion.

**Figure S4:**
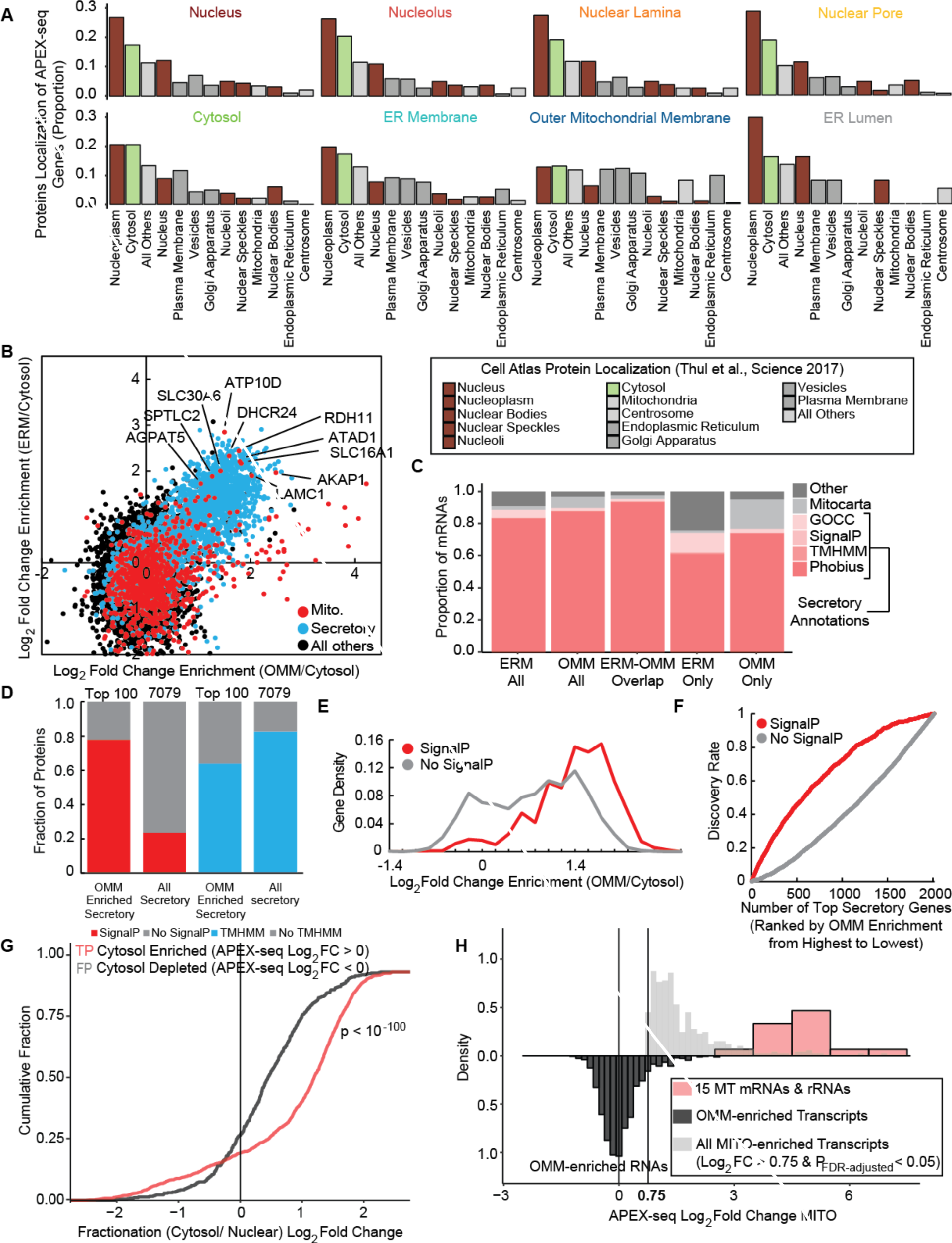
(A) Bar plots showing the protein localization of the transcripts enriched by APEX-seq. These numbers are based on the ~3250 genes examined in Figure 3 that have reliable protein localization data in the **Protein Cell Atlas** database. (B) Scatter plot comparing the OMM (x axis) and the ERM (y axis) APEX-seq log_2_fold-changes in HEK-293T cells. Genes are categorized as in Figure 6A. Gene names are shown for proteins known to be dual-localized to ER and mitochondria. (C) Of the mRNAs enriched by both OMM and ERM APEX-seq, more than 90% have secretory annotations. (D) Comparing the proportion of transcripts with SignalP or TMHMM prediction between top 100 secretory genes enriched by OMM APEX-seq in HEK 293T cells and all secretory genes (E) (F) Gene density distribution and ROC curve of secretory genes in OMM APEX-seq in HEK-293T cells under basal condition. Genes are ranked by their OMM APEX-seq log_2_ fold enrichment ratio from highest to lowest. (G) Cumulative distribution of transcripts in the nuclear/cytosolic fractionation data split into two groups based on cytosolic APEX-seq data. Genes enriched by cytosolic APEX-seq (log_2_ fold-change > 0, *p*_FDR-adjusted_ < 0.05) had much higher enrichment (*p* < 10^−100^, KS test), in the cytosolic fractionation data relative to genes depleted by cytosolic APEX-seq (log_2_ fold-change > 0, *p*_FDR-adjusted_ < 0.05). (H) Mitochondrial APEX-seq shows robust enrichment of the 15 MT rRNAs and mRNAs, and no enrichment of OMM-enriched RNAs. There are ~400 transcripts that have large positive fold changes (log_2_foldchange > 0.75, *p*_FDR-adjusted_ < 0.05).

**Figure S5:**
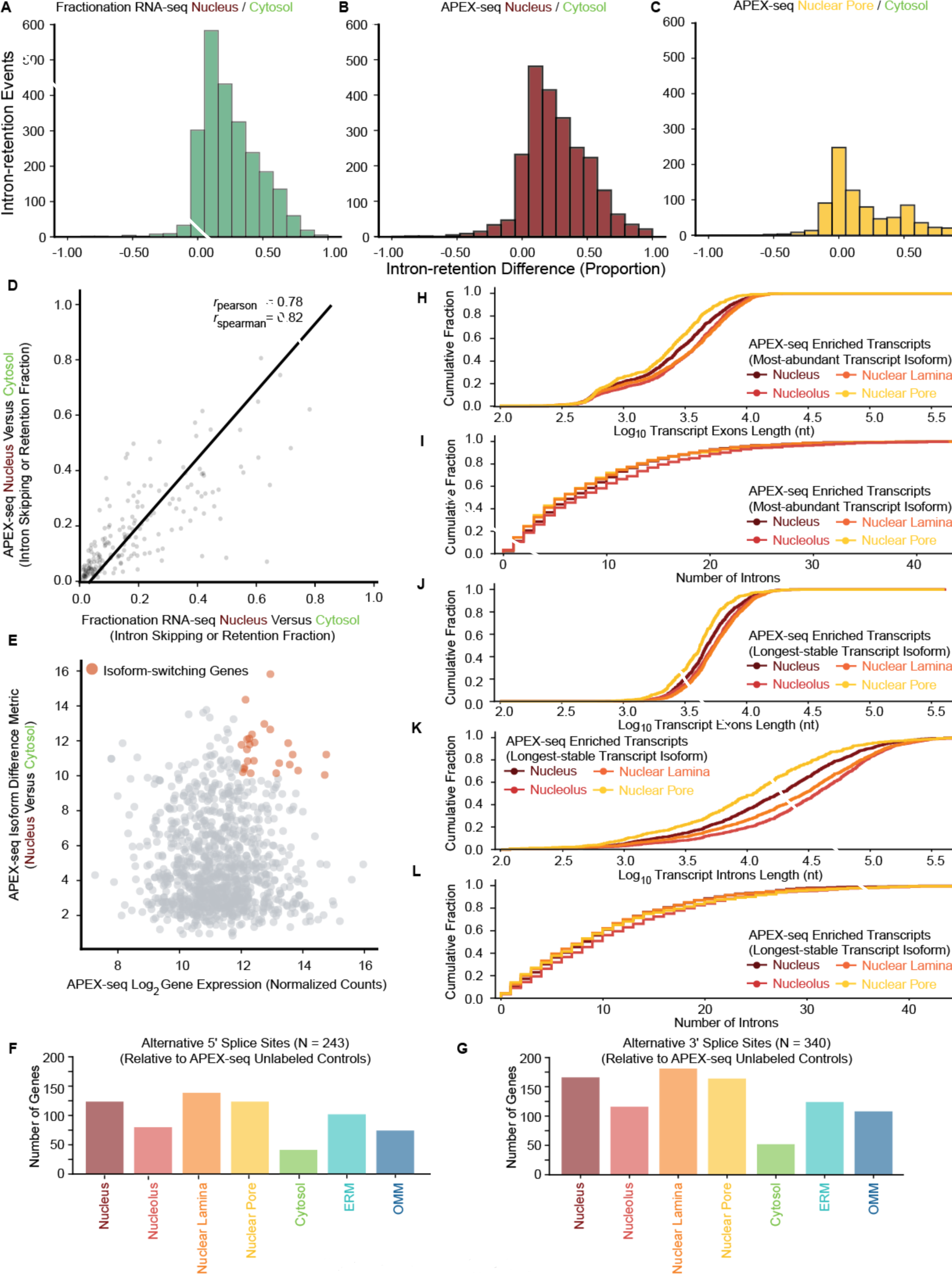
(A) (B) (C) Number of intron-retention events across APEX-seq enriched transcripts in the nucleus and nuclear pore, as well as fractionation RNA-seq. An intron-retention score was calculated based on how much the retained-intron transcript was obtained relative to the corresponding cytosol control, and was computed by taking the sum of the absolute values of the log2fold-change enrichments for the most cytosol-biased and most nuclear-biased transcript. (D) Scatter plot showing the high correlation between intron skipping or retention in nucleus APEX-seq (relative to cytosol APEX-seq) versus nuclear fractionation RNA-seq (relative to cytosol). Genes displaying no differential expression between the nucleus and cytosol, but with at least one transcript enriched in the nucleus and a different transcript enriched in the cytosol, were called as displaying isoform switching. (E) The genes shown in Figure 4E were identified by selecting for transcripts that are highly abundant and showed high-isoform switching scores. (F) (G) Barplots showing the number of genes showing alternative splice sites at (F) 5′ UTRs and 3′ UTRs in the APEX-seq samples, relative to unlabeled controls (FDR < 0.05). (H) (I) Cumulative distributions of the exon length, and number of isoforms for genes enriched by APEX-seq in the nuclear pore relative to other locations. We observe shorter transcripts at the nuclear pore relative to other locations. We see no significant difference in distribution across the locations. Here the transcript length was calculated by considering the most-abundance transcript isoform for each gene across all locations in the APEX-seq data. (J) (K) (L) Cumulative distribution of the introns length, exon lengths and number of introns for genes enriched by APEX-seq in the nuclear locations. Here the transcript length was calculated by considering the longest-stable transcript isoform for each gene.

**Figure S6:**
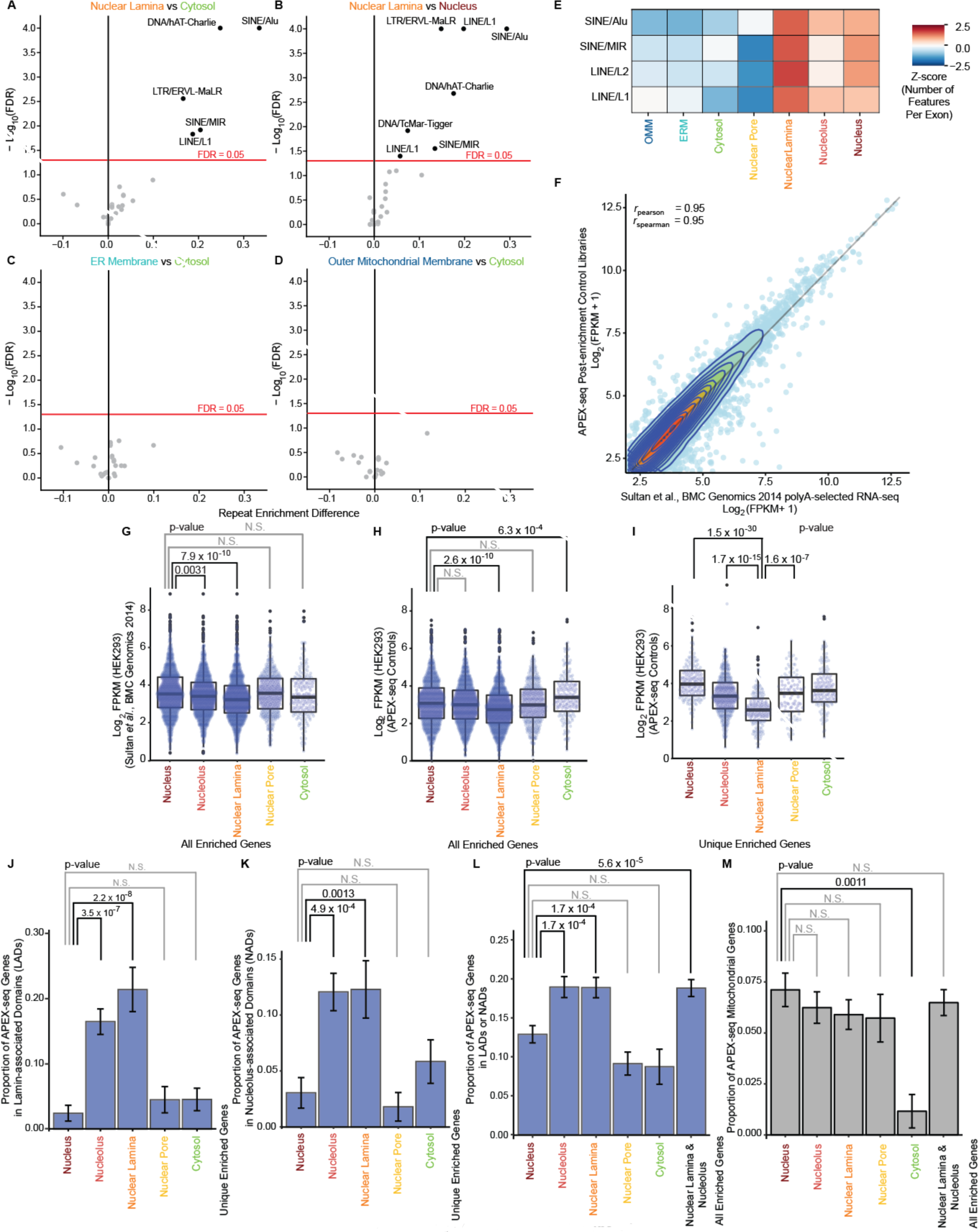
(A) (B) (C) (D)Using all the APEX-seq enriched genes as a background, the estimated FDR of finding the nuclear-lamina repeat motifs. (E) Heatmap showing the number of repeat motifs in exons of transcripts enriched by APEX-seq. Unlike in Figure 5A, this analysis considers all enriched genes, not just enriched genes unique to that location. We continue to see strong enrichment of these motifs in the nuclear lamina, but also in other nuclear locations relative to the cytosolic locations. (F) Scatter plot showing good correlation between post-enrichment APEX-seq control data and published polyA+ RNA-seq data from HEK293. APEX-seq control data was averaged from 18 controls generated from APEX2 constructs targeting 9 locations. (G) RNA-seq abundance of all genes (not just unique genes) enriched by APEX-seq. FPKM = fragments per kilobase per million reads. Abundance data from Sultan *et al*.(Sultan et al., 2014). (H) (I) Using post-enrichment APEX-seq control data we also obtain decreased abundance of nuclear-lamina enriched genes, both for all genes and more strikingly for unique genes. FPKM = fragments per kilobase per million reads. (I) (K) (L) Bar plots showing the proportion of genes found in lamina-associated domains or nucleolus associated domains or both.-values are from Fisher’s exact tests. (M) Control test examining the localization of mitochondrial genes, confirming no similar enrichment of genes in the nucleolus or nuclear-lamina transcriptomes. P-values are from Fisher’s exact tests.

**Figure S7:**
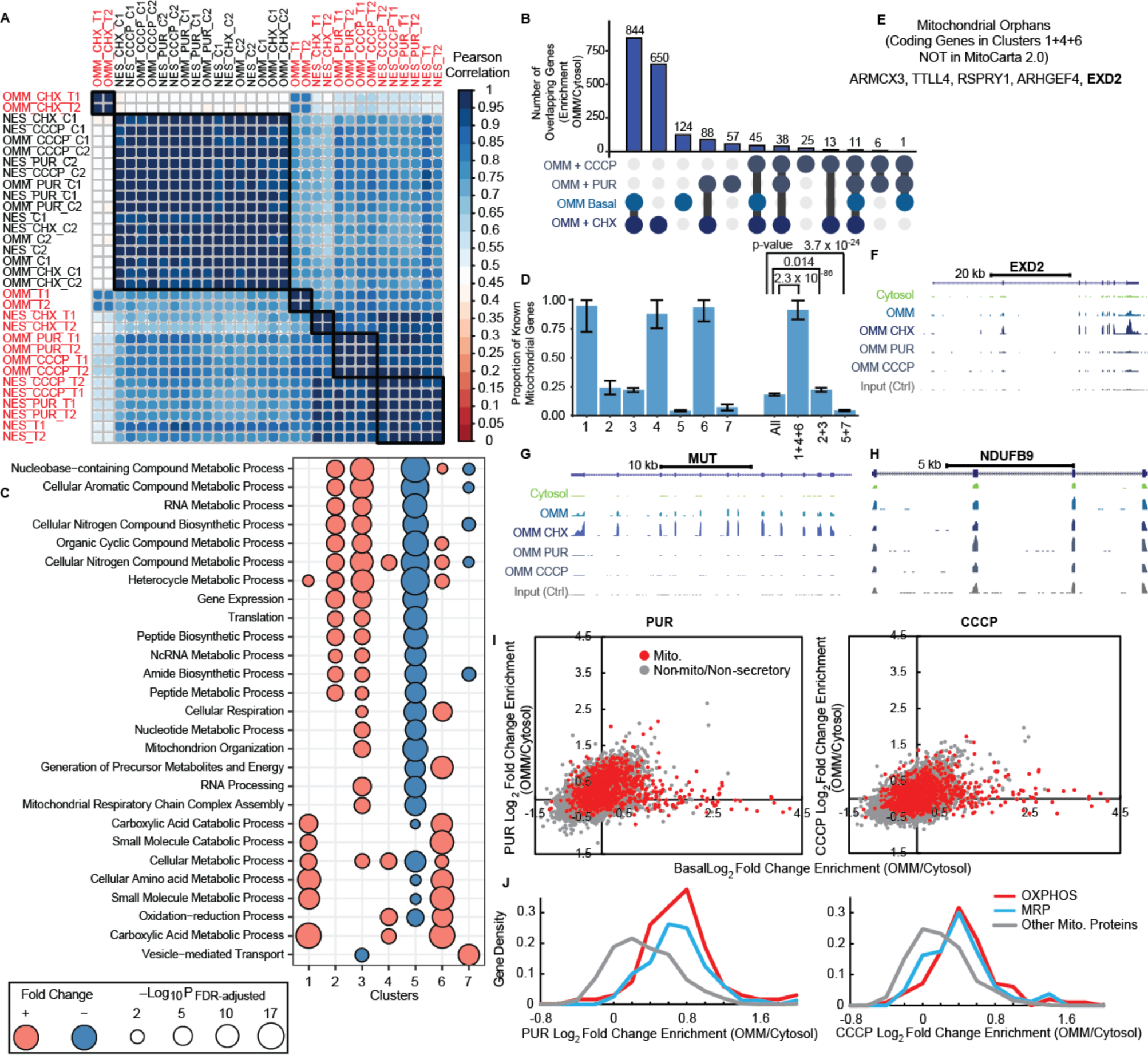
(A) Cluster map of the OMM perturbation experiments, along with the corresponding cytosolic background. All controls cluster together, while the cytosolic locations vary less across the different perturbation experiments relative to their OMM counterparts. OMM cycloheximide is the most different among these labeled libraries. (B) Plot showing the overlapping number of enriched genes in the different OMM perturbation experiments. All genes that were enriched in at least 1 of the four conditions are included. (C) Molecular function GO-terms based from clusters in Figure 6C. The clusters enriched in mitochondrial genes (1+4+6) show differences relative to clusters 2+3. (D) The proportion of known mitochondrial genes in the different clusters. Clusters 1+4+6 are highly enriched in mitochondrial genes, while clusters 5+7 are significantly depleted. (E) Examining the transcripts not annotated as mitochondrial in clusters 1+4+6 yields 12 transcripts, of which 7 are pseudogenes and 5 are mRNAs. Of these 5 mRNAs, further literature examination shows evidence for 1 coding for a protein localizing to the mitochondria (*TTLL4*(Thul et al., 2017)) and 2 localizing to the OMM (*ARMCX3*(Mou et al., 2009) and *EXD2*(Hensen et al., 2018)). (F) Genome tracks of *EXD2* from (E). (G) UCSC Browser tracks of a mitochondrial gene (MUT) show increased enrichment by OMM-APEX upon CHX treatment. (H) UCSC Browser tracks of an OXPHOS gene (*NDUFB6*) that show increased enrichment by OMM-APEX upon PUR/CCCP treatment. (I) ․Scatter plots of OMM APEX-seq log_2_ fold change in HEK-293T cells comparing the basal (x axis) and Puro (y axis, left)/CCCP (y axis, right) conditions. Genes are categorized as in Figure 6A. (J) Gene density distribution of OMM APEX-seq log_2_ fold-change in HEK-293T cells under Puro (left) or CCCP (right) condition. Genes are functionally classified according to Gene Ontology.

## REFERENCES

Bahar Halpern, K., Caspi, I., Lemze, D., Levy, M., Landen, S., Elinav, E., Ulitsky, I., and Itzkovitz, S. (2015). Nuclear Retention of mRNA in Mammalian Tissues. Cell Reports 13, 2653–2662.

Berkovits, B.D., and Mayr, C. (2015). Alternative 3’ UTRs act as scaffolds to regulate membrane protein localization. Nature 522, 363–367.

Bersuker, K., Peterson, C.W.H., To, M., Sahl, S.J., Savikhin, V., Grossman, E.A., Nomura, D.K., and Olzmann, J.A. (2018). A Proximity labeling strategy provides insights into the composition and dynamics of lipid droplet proteomes. Dev Cell 44, 97–112 e117.

Bertrand, E., Chartrand, P., Schaefer, M., Shenoy, S.M., Singer, R.H., and Long, R.M. (1998). Localization of ASH1 mRNA particles in living yeast. Mol Cell 2, 437–445.

Blobel, G. (1985). Gene gating: a hypothesis. Proc Natl Acad Sci USA 82, 8527–8529.

Boutz, P.L., Bhutkar, A., and Sharp, P.A. (2015). Detained introns are a novel, widespread class of post-transcriptionally spliced introns. Genes Dev 29, 63–80.

Brown, C.R., and Silver, P.A. (2007). Transcriptional regulation at the nuclear pore complex. Curr Opin Genet Dev 17, 100–106.

Buxbaum, A.R., Haimovich, G., and Singer, R.H. (2015). In the right place at the right time: visualizing and understanding mRNA localization. Nat Rev Mol Cell Biol 16, 95–109.

Cabili, M.N., Dunagin, M.C., McClanahan, P.D., Biaesch, A., Padovan-Merhar, O., Regev, A., Rinn, J.L., and Raj, A. (2015). Localization and abundance analysis of human lncRNAs at single cell and single-molecule resolution. Genome Biol 16, 20.

Calvo, S.E., Clauser, K.R., and Mootha, V.K. (2015). MitoCarta2.0: an updated inventory of mammalian mitochondrial proteins. Nuc Acids Res 44, D1251–D1257.

Carlevaro-Fita, J., Polidori T., Das M., Navarro C., and Johnson R. (2017). Ancient exapted transposable elements promote nuclear enrichment of long noncoding RNAs. bioRxiv

Chacinska, A., Koehler, C.M., Milenkovic, D., Lithgow, T., and Pfanner, N. (2009). Importing mitochondrial proteins: machineries and mechanisms. Cell 138, 628–644.

Chen, C.K., Blanco, M., Jackson, C., Aznauryan, E., Ollikainen, N., Surka, C., Chow, A., Cerase, A., McDonel, P., and Guttman, M. (2016). Xist recruits the X chromosome to the nuclear lamina to enable chromosome-wide silencing. Science 354, 468–472.

Chen, C.L., Hu, Y.H., Udeshi, N.D., Lau, T.Y., Wirtz-Peitz, F., He, L., Ting, A.Y., Carr, S.A., and Perrimon, N. (2015a). Proteomic mapping in live Drosophila tissues using an engineered ascorbate peroxidase. Proc Natl Acad Sci USA 112, 12093–12098.

Chen, K.H., Boettiger, A.N., Moffitt, J.R., Wang, S., and Zhuang, X. (2015b). Spatially resolved, highly multiplexed RNA profiling in single cells. Science 348, aaa6090–aaa6090.

Chen, K.W.Y.S.J. (2018). A statistical analysis on transcriptome sequences: the enrichment of Alu-element is associated with subcellular location. Biochem Biophys Res Comm 499, 397–402.

Chen, T., and van Steensel, B. (2017). Comprehensive analysis of nucleocytoplasmic dynamics of mRNA in Drosophila cells. PLoS Genetics 13, e1006929.

Chin, A., and Lecuyer, E. (2017). RNA localization: making its way to the center stage. Biochim Biophys Acta 1861, 2956–2970.

Couvillion, M.T., Soto, I.C., Shipkovenska, G., and Churchman, L.S. (2016). Synchronized mitochondrial and cytosolic translation programs. Nature 533, 499–503.

Dai, J., Sloat, A.L., Wright, M.W., and Manderville, R.A. (2005). Role of phenoxyl radicals in DNA adduction by chlorophenol xenobiotics following peroxidase activation. Chem Res Toxicology 18, 771–779.

Dai, J., Wright, M.W., and Manderville, R.A. (2003a). Ochratoxin A forms a carbon-bonded C8-deoxyguanosine nucleoside adduct: implications for C8 reactivity by a phenolic radical. J Am Chem Soc 125, 3716–3717.

Dai, J., Wright, M.W., and Manderville, R.A. (2003b). An oxygen-bonded C8-deoxyguanosine nucleoside adduct of pentachlorophenol by peroxidase activation: evidence for ambident C8 reactivity by phenoxyl radicals. Chem Res Toxicology 16, 817–821.

de Koning, A.P., Gu, W., Castoe, T.A., Batzer, M.A., and Pollock, D.D. (2011). Repetitive elements may comprise over two-thirds of the human genome. PLoS Genet 7, e1002384.

Dekker, J., Belmont, A.S., Guttman, M., Leshyk, V.O., Lis, J.T., Lomvardas, S., Mirny, L.A., O’Shea, C.C., Park, P.J., Ren, B., et al. (2017). The 4D nucleome project. Nature 549, 219–226.

Delaleau, M., and Borden, K.L. (2015). Multiple Export Mechanisms for mRNAs. Cells 4, 452–473.

Dillinger, S., Straub, T., and Nemeth, A. (2017). Nucleolus association of chromosomal domains is largely maintained in cellular senescence despite massive nuclear reorganisation. PLoS One 12, e0178821.

Djebali, S., Davis, C.A., Merkel, A., Dobin, A., Lassmann, T., Mortazavi, A., Tanzer, A., Lagarde, J., Lin, W., Schlesinger, F., et al. (2012). Landscape of transcription in human cells. Nature 489, 101–108.

Fasken, M.B., and Corbett, A.H. (2009). Mechanisms of nuclear mRNA quality control. RNA Biol 6, 237–241.

Femino, A.M., Fay, F.S., Fogarty, K., and Singer, R.H. (1998). Visualization of single RNA transcripts in situ. Science 280, 585–590.

Fox, C.H., Johnson, F.B., Whiting, J., and Roller, P.P. (1985). Formaldehyde fixation. J Histochem Cytochem 33, 845–853.

Friedman, J.R., Lackner, L.L., West, M., DiBenedetto, J.R., Nunnari, J., and Voeltz, G.K. (2011). ER tubules mark sites of mitochondrial division. Science 334, 358–362.

Giacomello, M., and Pellegrini, L. (2016). The coming of age of the mitochondria–ER contact: a matter of thickness. Cell Death Diff 23, 1417–1427.

Gold, V.A., Chroscicki, P., Bragoszewski, P., and Chacinska, A. (2017). Visualization of cytosolic ribosomes on the surface of mitochondria by electron cryo-tomography. EMBO Rep 18, 1786–1800.

Grunwald, D., and Singer, R.H. (2010). In vivo imaging of labelled endogenous beta-actin mRNA during nucleocytoplasmic transport. Nature 467, 604–607.

Grunwald, D., Singer, R.H., and Rout, M. (2011). Nuclear export dynamics of RNA-protein complexes. Nature 475, 333–341.

Guelen, L., Pagie, L., Brasset, E., Meuleman, W., Faza, M.B., Talhout, W., Eussen, B.H., de Klein, A., Wessels, L., de Laat, W., et al. (2008). Domain organization of human chromosomes revealed by mapping of nuclear lamina interactions. Nature 453, 948–951.

Gupta, R., Somyajit, K., Narita, T., Maskey, E., Stanlie, A., Kremer, M., Typas, D., Lammers, M., Mailand, N., Nussenzweig, A., et al. (2018). DNA Repair Network Analysis Reveals Shieldin as a Key Regulator of NHEJ and PARP Inhibitor Sensitivity. Cell 173, 972–988 e923.

Han, S., Udeshi, N.D., Deerinck, T.J., Svinkina, T., Ellisman, M.H., Carr, S.A., and Ting, A.Y. (2017). Proximity biotinylation as a method for mapping proteins associated with mtDNA in living cells. Cell Chem Biol 24, 404–414.

Hensen, F., Moretton, A., van Esveld, S., Farge, G., and Spelbrink, J.N. (2018). The mitochondrial outer-membrane location of the EXD2 exonuclease contradicts its direct role in nuclear DNA repair. Sci Rep 8, 5368.

Hubbard, T., Barker, D., Birney, E., Cameron, G., Chen, Y., Clark, L., Cox, T., Cuff, J., Curwen, V., Down, T., et al. (2002). The Ensembl genome database project. Nuc Acids Res 30, 38–41.

Hubley, R., Finn, R.D., Clements, J., Eddy, S.R., Jones, T.A., Bao, W., Smit, A.F., and Wheeler, T.J. (2016). The Dfam database of repetitive DNA families. Nuc Acids Res 44, D81–89.

Hung, V., Lam, S.S., Udeshi, N.D., Svinkina, T., Guzman, G., Mootha, V.K., Carr, S.A., and Ting, A.Y. (2017). Proteomic mapping of cytosol-facing outer mitochondrial and ER membranes in living human cells by proximity biotinylation. ELife 6.

Hung, V., Udeshi, N.D., Lam, S.S., Loh, K.H., Cox, K.J., Pedram, K., Carr, S.A., and Ting, A.Y. (2016). Spatially resolved proteomic mapping in living cells with the engineered peroxidase APEX2. Nat Protoc 11, 456–475.

Hung, V., Zou, P., Rhee, H.-W., Udeshi, Namrata D., Cracan, V., Svinkina, T., Carr, Steven A., Mootha, Vamsi K., and Ting, Alice Y. (2014). Proteomic mapping of the human mitochondrial intermembrane space in live cells via ratiometric APEX tagging. Mol Cell 55, 332–341.

Hwang, J., and Espenshade, P.J. (2016). Proximity-dependent biotin labelling in yeast using the engineered ascorbate peroxidase APEX2. Biochem J 473, 2463–2469.

Ichiyanagi, K. (2013). Epigenetic regulation of transcription and possible functions of mammalian short interspersed elements, SINEs. Genes Genet Syst 88, 19–29.

Ingolia, N.T., Ghaemmaghami, S., Newman, J.R., and Weissman, J.S. (2009). Genome-wide analysis in vivo of translation with nucleotide resolution using ribosome profiling. Science 324, 218–223.

Jan, C.H., Williams, C.C., and Weissman, J.S. (2014). Principles of ER cotranslational translocation revealed by proximity-specific ribosome profiling. Science 346, 1257521–1257521.

Kaewsapsak, P., Shechner, D.M., Mallard, W., Rinn, J.L., and Ting, A.Y. (2017). Live-cell mapping of organelle-associated RNAs via proximity biotinylation combined with protein-RNA crosslinking. Elife 6.

Kall, L., Krogh, A., and Sonnhammer, E.L. (2004). A combined transmembrane topology and signal peptide prediction method. J Mol Biol 338, 1027–1036.

Katahira, J. (2015). Nuclear export of messenger RNA. Genes 6, 163–184.

Kellems, R.E., Allison, V.F., and Butow, R.A. (1974). Cytoplasmic type 80 S ribosomes associated with yeast mitochondria. II. Evidence for the association of cytoplasmic ribosomes with the outer mitochondrial membrane in situ. J Biol Chem 249, 3297–3303.

Kellems, R.E., Allison, V.F., and Butow, R.A. (1975). Cytoplasmic type 80S ribosomes associated with yeast mitochondria. IV. Attachment of ribosomes to the outer membrane of isolated mitochondria. J Cell Biol 65, 1–14.

Kent, W.J., Sugnet, C.W., Furey, T.S., Roskin, K.M., Pringle, T.H., Zahler, A.M., and Haussler, D. (2002). The human genome browser at UCSC. Genome Res 12, 996–1006.

Khong, A., Matheny, T., Jain, S., Mitchell, S.F., Wheeler, J.R., and Parker, R. (2017). The stress granule transcriptome reveals principles of mRNA accumulation in stress granules. Mol Cell 68, 808–820 e805.

Kim, S.J., Fernandez-Martinez, J., Nudelman, I., Shi, Y., Zhang, W., Raveh, B., Herricks, T., Slaughter, B.D., Hogan, J.A., Upla, P., et al. (2018). Integrative structure and functional anatomy of a nuclear pore complex. Nature 555, 475–482.

Kornmann, B., Currie, E., Collins, S.R., Schuldiner, M., Nunnari, J., Weissman, J.S., and Walter, P. (2009). An ER-mitochondria tethering complex revealed by a synthetic biology screen. Science 325, 477–481.

Krogh, A., Larsson, B., von Heijne, G., and Sonnhammer, E.L. (2001). Predicting transmembrane protein topology with a hidden Markov model: application to complete genomes. J Mol Biol 305, 567–580.

Lam, S.S., Martell, J.D., Kamer, K.J., Deerinck, T.J., Ellisman, M.H., Mootha, V.K., and Ting, A.Y. (2014). Directed evolution of APEX2 for electron microscopy and proximity labeling. Nat Methods 12, 51–54.

Lee, J.H., Daugharthy, E.R., Scheiman, J., Kalhor, R., Yang, J.L., Ferrante, T.C., Terry, R., Jeanty, S.S.F., Li, C., Amamoto, R., et al. (2014). Highly multiplexed subcellular RNA sequencing in situ. Science 343, 1360–1363.

Lesnik, C., Golani-Armon, A., and Arava, Y. (2015). Localized translation near the mitochondrial outer membrane: an update. RNA Biol 12, 801–809.

Linden, A. (2006). Measuring diagnostic and predictive accuracy in disease management: an introduction to receiver operating characteristic (ROC) analysis. J Eval Clin Pract 12, 132–139.

Liu, N., Lee, C.H., Swigut, T., Grow, E., Gu, B., Bassik, M.C., and Wysocka, J. (2018). Selective silencing of euchromatic L1s revealed by genome-wide screens for L1 regulators. Nature 553, 228–232.

Lobingier, B.T., Hüttenhain, R., Eichel, K., Miller, K.B., Ting, A.Y., von Zastrow, M., and Krogan, N.J. (2017). An approach to spatiotemporally resolve protein interaction networks in living cells. Cell 169, 350–360.e312.

Loh, K.H., Stawski, P.S., Draycott, A.S., Udeshi, N.D., Lehrman, E.K., Wilton, D.K., Svinkina, T., Deerinck, T.J., Ellisman, M.H., Stevens, B., et al. (2016). Proteomic analysis of unbounded cellular compartments: synaptic clefts. Cell 166, 1295–1307.e1221.

Love, M.I., Huber, W., and Anders, S. (2014). Moderated estimation of fold change and dispersion for RNA-seq data with DESeq2. Genome Biol 15.

Lubelsky, Y., and Ulitsky, I. (2018). Sequences enriched in Alu repeats drive nuclear localization of long RNAs in human cells. Nature 555, 107–111.

Ma, J., Liu, Z., Michelotti, N., Pitchiaya, S., Veerapaneni, R., Androsavich, J.R., Walter, N.G., and Yang, W. (2013). High-resolution three-dimensional mapping of mRNA export through the nuclear pore. Nat Comm 4, 2414.

Markmiller, S., Soltanieh, S., Server, K.L., Mak, R., Jin, W., Fang, M.Y., Luo, E.C., Krach, F., Yang, D., Sen, A., et al. (2018). Context-dependent and disease-specific diversity in protein interactions within stress granules. Cell 172, 590–604 e513.

McCloskey, A., Taniguchi, I., Shinmyozu, K., and Ohno, M. (2012). hnRNP C tetramer measures RNA length to classify RNA polymerase II transcripts for export. Science 335, 1643–1646.

Mercer, Tim R., Neph, S., Dinger, Marcel E., Crawford, J., Smith, Martin A., Shearwood, A. Marie J., Haugen, E., Bracken, Cameron P., Rackham, O., Stamatoyannopoulos, John A., et al. (2011). The human mitochondrial transcriptome. Cell 146, 645–658.

Meyer, K.D., Saletore, Y., Zumbo, P., Elemento, O., Mason, C.E., and Jaffrey, S.R. (2012). Comprehensive analysis of mRNA methylation reveals enrichment in 3’ UTRs and near stop codons. Cell 149, 1635–1646.

Mortazavi, A., Williams, B.A., McCue, K., Schaeffer, L., and Wold, B. (2008). Mapping and quantifying mammalian transcriptomes by RNA-Seq. Nat Methods 5, 621–628.

Mortensen, A., and Skibsted, L.H. (1997). Importance of carotenoid structure in radical scavenging reactions. J Agr Food Chem 45, 2970–2977.

Mou, Z., Tapper, A.R., and Gardner, P.D. (2009). The armadillo repeat-containing protein, ARMCX3, physically and functionally interacts with the developmental regulatory factor Sox10. J Biol Chem 284, 13629–13640.

Padeken, J., Zeller, P., and Gasser, S.M. (2015). Repeat DNA in genome organization and stability. Curr Opin Genet Dev 31, 12–19.

Paek, J., Kalocsay, M., Staus, D.P., Wingler, L., Pascolutti, R., Paulo, J.A., Gygi, S.P., and Kruse, A.C. (2017). Multidimensional tracking of GPCR signaling via peroxidase-catalyzed proximity labeling. Cell 169, 338–349.e311.

Pandey, R.R., Homolka, D., Chen, K.M., Sachidanandam, R., Fauvarque, M.O., and Pillai, R.S. (2017). Recruitment of Armitage and Yb to a transcript triggers its phased processing into primary piRNAs in Drosophila ovaries. PLoS Genet 13, e1006956.

Patel, A., Lee, H.O., Jawerth, L., Maharana, S., Jahnel, M., Hein, M.Y., Stoynov, S., Mahamid, J., Saha, S., Franzmann, T.M., et al. (2015). A liquid-to-solid phase transition of the ALS Protein FUS accelerated by disease mutation. Cell 162, 1066–1077.

Penny, G.D., Kay, G.F., Sheardown, S.A., Rastan, S., and Brockdorff, N. (1996). Requirement for Xist in X chromosome inactivation. Nature 379, 131–137.

Petersen, T.N., Brunak, S., von Heijne, G., and Nielsen, H. (2011). SignalP 4.0: discriminating signal peptides from transmembrane regions. Nat Methods 8, 785–786.

Reed, R., and Cheng, H. (2005). TREX, SR proteins and export of mRNA. Curr Opin Cell Biol 17, 269–273.

Reid, D.W., and Nicchitta, C.V. (2012). Primary role for endoplasmic reticulum-bound ribosomes in cellular translation identified by ribosome profiling. J Biol Chem 287, 5518–5527.

Reinke, A.W., Mak, R., Troemel, E.R., and Bennett, E.J. (2017). In vivo mapping of tissue- and subcellular-specific proteomes in Caenorhabditis elegans. Sci Adv 3, e1602426.

Rhee, H.W., Zou, P., Udeshi, N.D., Martell, J.D., Mootha, V.K., Carr, S.A., and Ting, A.Y. (2013). Proteomic mapping of mitochondria in living cells via spatially restricted enzymatic tagging. Science 339, 1328–1331.

Richter-Dennerlein, R., Oeljeklaus, S., Lorenzi, I., Ronsor, C., Bareth, B., Schendzielorz, A.B., Wang, C., Warscheid, B., Rehling, P., and Dennerlein, S. (2016). Mitochondrial protein synthesis adapts to influx of nuclear-encoded protein. Cell 167, 471–483 e410.

Roundtree, I.A., Luo, G.Z., Zhang, Z., Wang, X., Zhou, T., Cui, Y., Sha, J., Huang, X., Guerrero, L., Xie, P., et al. (2017). YTHDC1 mediates nuclear export of N(6)-methyladenosine methylated mRNAs. Elife 6.

Sadowski, P.G., Groen, A.J., Dupree, P., and Lilley, K.S. (2008). Sub-cellular localization of membrane proteins. Proteomics 8, 3991–4011.

Schnell, U., Dijk, F., Sjollema, K.A., and Giepmans, B.N.G. (2012). Immunolabeling artifacts and the need for live-cell imaging. Nat Methods 9, 152–158.

Shah, S., Lubeck, E., Zhou, W., and Cai, L. (2016). In situ transcription profiling of single cells reveals spatial organization of cells in the mouse hippocampus. Neuron 92, 342–357.

Shukla, C.J., McCorkindale, A.L., Gerhardinger, C., Korthauer, K.D., Cabili, M.N., Shechner, D.M., Irizarry, R.A., Maass, P.G., and Rinn, J.L. (2018). High-throughput identification of RNA nuclear enrichment sequences. EMBO J 37.

Singh, G., Kucukural, A., Cenik, C., Leszyk, J.D., Shaffer, S.A., Weng, Z., and Moore, M.J. (2012). The cellular EJC interactome reveals higher-order mRNP structure and an EJC-SR protein nexus. Cell 151, 750–764.

Smart, A.C., Margolis, C.A., Pimentel, H., He, M.X., Miao, D., Adeegbe, D., Fugmann, T., Wong, K.K., and Van Allen, E.M. (2018). Intron retention is a source of neoepitopes in cancer. Nat Biotech.

Spitale, R.C., Flynn, R.A., Zhang, Q.C., Crisalli, P., Lee, B., Jung, J.-W., Kuchelmeister, H.Y., Batista, P.J., Torre, E.A., Kool, E.T., et al. (2015). Structural imprints in vivo decode RNA regulatory mechanisms. Nature 519, 486–490.

Sultan, M., Amstislavskiy, V., Risch, T., Schuette, M., Dokel, S., Ralser, M., Balzereit, D., Lehrach, H., and Yaspo, M.L. (2014). Influence of RNA extraction methods and library selection schemes on RNA-seq data. BMC Genomics 15, 675.

Thul, P.J., Akesson, L., Wiking, M., Mahdessian, D., Geladaki, A., Ait Blal, H., Alm, T., Asplund, A., Bjork, L., Breckels, L.M., et al. (2017). A subcellular map of the human proteome. Science 356.

Valm, A.M., Cohen, S., Legant, W.R., Melunis, J., Hershberg, U., Wait, E., Cohen, A.R., Davidson, M.W., Betzig, E., and Lippincott-Schwartz, J. (2017). Applying systems-level spectral imaging and analysis to reveal the organelle interactome. Nature 546, 162–167.

van Koningsbruggen, S., Gierlinski, M., Schofield, P., Martin, D., Barton, G.J., Ariyurek, Y., den Dunnen, J.T., and Lamond, A.I. (2010). High-resolution whole-genome sequencing reveals that specific chromatin domains from most human chromosomes associate with nucleoli. Mol Biol Cell 21, 3735–3748.

Van Nostrand, E.L., Pratt, G.A., Shishkin, A.A., Gelboin-Burkhart, C., Fang, M.Y., Sundararaman, B., Blue, S.M., Nguyen, T.B., Surka, C., Elkins, K., et al. (2016). Robust transcriptome-wide discovery of RNA-binding protein binding sites with enhanced CLIP (eCLIP). Nat Methods 13, 508–514.

van Steensel, B., and Belmont, A.S. (2017). Lamina-associated domains: links with chromosome architecture, heterochromatin, and gene repression. Cell 169, 780–791.

Walther, T.C., Fornerod, M., Pickersgill, H., Goldberg, M., Allen, T.D., and Mattaj, I.W. (2001). The nucleoporin Nup153 is required for nuclear pore basket formation, nuclear pore complex anchoring and import of a subset of nuclear proteins. EMBO J 20, 5703–5714.

Wang, X., Allen, W.E., Wright, M.A., Sylwestrak, E.L., Samusik, N., Vesuna, S., Evans, K., Liu, C., Ramakrishnan, C., Liu, J., et al. (2018). Three-dimensional intact-tissue sequencing of single-cell transcriptional states. Science 361.

Weil, T.T., Parton, R.M., and Davis, I. (2010). Making the message clear: visualizing mRNA localization. Trends Cell Biol 20, 380–390.

Wheeler, T.J., Clements, J., Eddy, S.R., Hubley, R., Jones, T.A., Jurka, J., Smit, A.F., and Finn, R.D. (2013). Dfam: a database of repetitive DNA based on profile hidden Markov models. Nuc Acids Res 41, D70–82.

Wickramasinghe, V.O., and Laskey, R.A. (2015). Control of mammalian gene expression by selective mRNA export. Nat Rev Mol Cell Biol 16, 431–442.

Williams, C.C., Jan, C.H., and Weissman, J.S. (2014). Targeting and plasticity of mitochondrial proteins revealed by proximity-specific ribosome profiling. Science 346, 748–751.

Wishart, J.F., and Rao, B.S.M. (2010). Recent trends in radiation chemistry (World Scientific).

